# Cross-tissue analysis of allelic X-chromosome inactivation ratios resolves features of human development

**DOI:** 10.1101/2021.09.13.460075

**Authors:** Jonathan M. Werner, Sara Ballouz, John Hover, Jesse Gillis

## Abstract

X-chromosome inactivation (XCI) is a random, permanent, and developmentally early epigenetic event that occurs during mammalian embryogenesis. We harness these features of XCI to investigate characteristics of early lineage specification events during human development. We initially assess the consistency of X-inactivation and establish a robust set of XCI-escape genes. By analyzing variance in XCI ratios across tissues and individuals, we find that XCI is completed prior to tissue specification and at a time when 6-16 cells are fated for all tissue lineages. Additionally, we exploit tissue specific variability to characterize the number of cells present at the time of each tissue’s lineage commitment, ranging from approximately 20 cells in liver and whole blood tissues to 80 cells in brain tissues. By investigating variance of XCI ratios using adult tissue, we resolve key features of human development otherwise difficult to ascertain experimentally and develop scalable methods easily applicable to future data.

## Introduction

Every cell within female mammalian embryos undergoes the process of X-chromosome inactivation (XCI), which silences expression from a single randomly chosen X-allele via epigenetic mechanisms^1,2,3^. The random choice of which allele to inactivate occurs early in development and is permanent thereafter with the inactivated allele propagated through each cell’s developmental lineage^4^. As a result, adult females exhibit mosaic X-linked allelic expression throughout every tissue within the body, an enduring phenotypic consequence of an early embryonic milestone. The random, permanent, and developmentally early nature of XCI positions the whole-body mosaicism of X-linked allelic expression as a lineage marker reaching back to the earliest embryonic stages^5,6^. Careful analysis of X-linked allelic expression across individuals and tissues can thus reveal whole-body lineage relationships stemming from some of the first lineage decisions made during embryogenesis^6,7,8,9^.

While the probability of inactivation is equal between the X-alleles in humans, variation in XCI allelic ratios across individuals is a salient feature of XCI. Deviation from the expected XCI allelic ratio of 0.5 can arise through various mechanisms^10,11,12^ with the most basic being the inherent stochasticity of the initial choice of allelic inactivation^13^. The variability of the initial XCI ratio within the embryo is directly linked to the number of cells present during inactivation where smaller cell numbers result in increased variability of XCI ratios^6^. In fact, one can estimate the number of cells present at the time of inactivation by analyzing the variance of XCI ratios across a population. Several studies using this approach^13,14^, as well as studies using *in vitro* embryonic models^15,16,17^, have estimated that XCI occurs in a small stem cell pool within the human embryo with estimates as little as 8 cells. The combination of the random nature and small pool of cells present during XCI imparts an ever-present basal-level of variability in XCI ratios within adult human populations.

The stability of XCI down lineages means that minor cell sampling variation can be used as a marker for any process involving selection of a set of cells, i.e., lineage specification^6,7^. While growing evidence indicates XCI is initiated early^15,16,17^, the timing of XCI completion is unclear^18^ and has important implications for the variance in XCI ratios across early lineages. Germ layer specification is the first lineage decision made for all future embryonic tissues and occurs during post-implantation embryonic development^19^, a similar timeframe to XCI. If XCI is completed before germ layer specification each germ layer would be specified from the same pool of cells with a set XCI ratio (Fig. 1a). The germ layer specific XCI ratio would be dependent on the initial XCI ratio resulting in shared XCI ratios across germ layers (Fig. 1a). In contrast, if XCI is completed after germ layer specification, the germ layer specific XCI ratio is set independently and is not expected to be shared across the different germ layers (Fig. 1b). Consequently, comparing XCI ratios for tissues within either the same or different germ layer lineages can reveal the temporal ordering of XCI and germ layer specification. Extending to tissue-specific lineages, variability in the number of cells fated for individual tissues will generate stereotypic deviations in XCI ratios due to sampling, potentially allowing estimates of cell numbers for tissue-specific lineage commitment.

**Figure 1.**
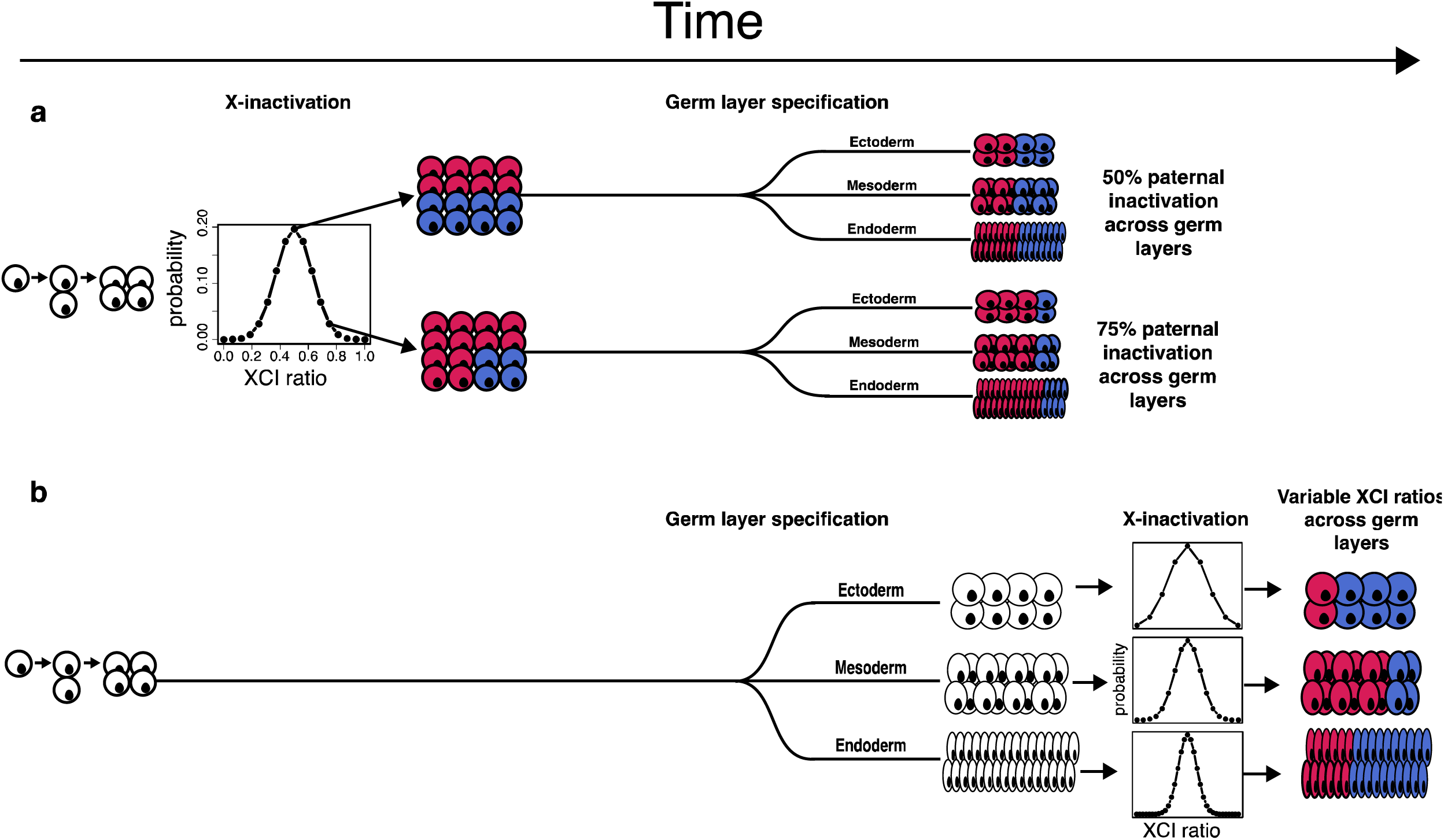
Timing of XCI determines lineage-specific XCI ratio probability. **a**, Schematic representing completed XCI before germ layer specification. Each germ layer inherits the same randomly determined XCI ratio set prior to germ layer lineage specification. The probability distribution of XCI is determined by the number of cells present during inactivation. **b**, Schematic representing completed XCI after germ layer specification. The XCI ratio for each germ layer is set independent of one another, together along with variation in cell numbers fated for each germ layer results in variable XCI ratios across the germ layer lineages.

In this study, we develop an approach to determine the tissue XCI ratio from unphased bulk RNA-sequencing data allowing us to assess XCI ratios from any publicly available RNA-sequencing dataset. Utilizing the tissue sampling scheme of the Genotype-Tissue Expression (GTEx v8) project^20^, we analyze XCI ratios for 49 tissues both within and across individuals for 311 female donors (Supp. Fig. 1). We establish that XCI ratios are shared for tissues both within and across germ layers demonstrating that XCI is completed before any significant lineage decisions are made for embryonic tissues. Additionally, we extend population-level modeling of variance in XCI ratios to all well-powered tissues, deriving estimates for the number of cells present at the time of XCI and tissue-specific lineage commitment. By providing cell counts, temporal ordering of lineage events, and lineage relationships across tissues, capturing the statistical commonalities that underlie the inherently stochastic nature of XCI is a powerful approach for resolving questions of early developmental lineage specification.

## Results

### The folded-normal model accurately estimates XCI ratios from unphased data

A practical consequence of bulk RNA-sequencing is that the XCI ratio of a tissue can be estimated from the direction and magnitude of X-linked allele-specific expression. For a tissue with 75% of cells carrying an active maternal X-allele, approximately 75% of RNA-sequencing reads for heterozygous loci are expected to align to the maternal X-allele (Fig. 2a). However, allelic expression for any given gene is affected by a variety of factors both biological (e.g., eQTLs) and technical (e.g., read sampling). To derive robust estimates, we aggregate allelic expression ratios across well-powered intra-genic heterozygous SNPs for a given tissue, providing a chromosome wide-estimate of the tissue XCI ratio (Fig, 2a).

**Figure 2.**
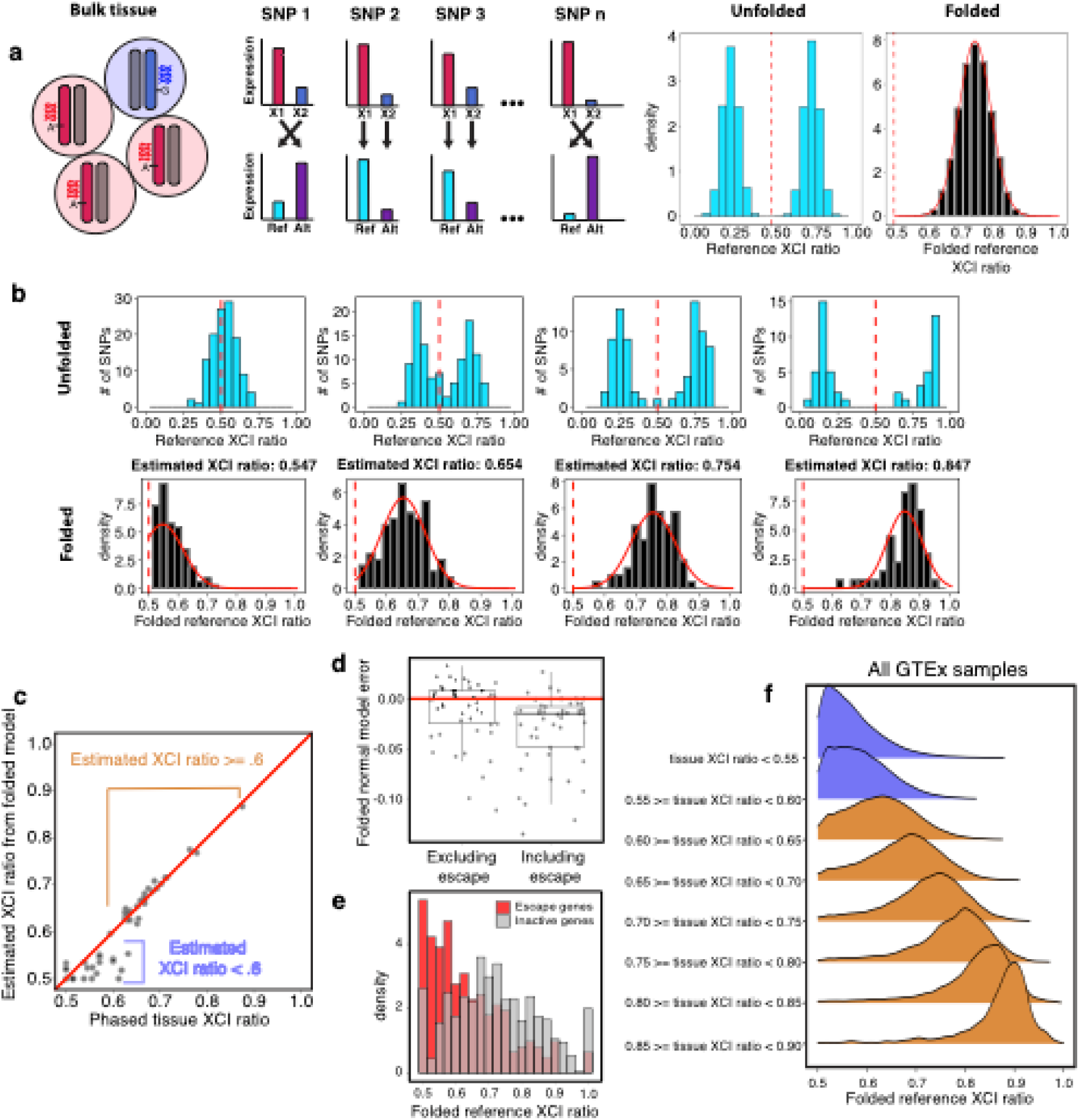
The folded-normal model accurately estimates XCI ratios from unphased bulk RNA-sequencing data. **a**, Schematic demonstrating how allelic expression of heterozygous SNPs reflect the XCI ratio of bulk tissue samples. Aligning expression data to a reference genome scrambles the parental haplotypes and reference allelic expression ratios will represent both the expected maternal and paternal allelic expression ratios given the XCI ratio of the tissue. Folding the reference allelic expression ratios captures the magnitude of the tissue XCI ratio. **b**, Distributions of reference allelic expression ratios for identified heterozygous SNPs across tissue samples exhibiting a range of bulk XCI ratios. Both the unfolded (top row) and folded distributions with the fitted folded normal model (bottom row) are shown. **c**, For the EN-TEx tissue samples, the phased median gene XCI ratio is plotted against the unphased XCI ratio estimate from the folded normal model. The folded normal model produces near identical XCI ratio estimates for samples with XCI ratios greater than or equal to 0.60. **d**, Deviation of the folded normal model from the phased median gene XCI ratio when excluding or including known escape genes. **e**, Aggregated folded reference allelic expression distributions for known escape and inactive genes in EN-TEx tissues with XCI ratios >= 0.70. Balanced allelic expression in opposition to the XCI ratio of the tissue most likely explains the increased underestimation of XCI ratios from the folded normal model when including escape genes. **f**, Aggregated gene-level folded reference allelic expression ratio distributions for all GTEx samples grouped by their estimated tissue XCI ratios (withholding known escape genes). The long tails of the distributions towards balanced biallelic expression are indicative of novel escape genes.

When aligned to a reference genome, reference alleles will be composed of both maternal and paternal alleles for a given sample. It follows that reference allelic expression ratios represent the expected expression ratios from both the maternal and paternal alleles given the XCI ratio of the tissue (Fig. 2a). To account for this, folding the reference allelic expression ratios about 0.5 aggregates the imbalanced allelic expression within the tissue across the two alleles, enabling the magnitude of the XCI ratio to be estimated from unphased expression data by fitting a folded distribution^21,22^ (see methods, Fig. 2a, b).

To assess the accuracy of the folded-normal model in estimating XCI ratios, we test our approach with phased bulk RNA-sequencing data from the EN-TEx^23,24^ consortium, a total of 49 tissue samples from 2 female donors spanning 26 different tissues. Comparing the unphased estimates derived with the folded normal model to the phased median allelic expression per sample, we find nearly perfect XCI ratio estimate correspondence for ratios greater than 0.6 (Fig. 2c). For samples skewed closer to the folding point of 0.5, model misspecification of the underlying distribution makes the estimate overconservative.

Our approach for estimating XCI ratios exploits the fact that aggregating allelic expression across numerous heterozygous loci averages mechanisms outside of XCI that may impact the allelic expression of X-linked genes. An intriguing and well-documented example of such a mechanism is escape from inactivation where a gene is biallelically expressed from both the active and inactive X-alleles within a cell^25^. Escape from inactivation is relatively common with between 15-30% of genes on the X-chromosome exhibiting evidence for escape^25,26^. While we exclude known escape genes^25^ from our folded-normal XCI ratio estimates, it is very likely unannotated escape genes are present within the data and may impact our XCI ratio estimates. To identify the impact of escape on our XCI ratio estimates, we derive folded-normal XCI ratio estimates while either excluding or including known escape genes and then compare these to the phased XCI ratio of tissues excluding the known escape genes (Fig. 2d). Including known escape genes results in an increased bias for underestimated XCI ratios with the folded-normal model (Fig. 2d). By comparing allelic ratios of known escape genes to all other genes in EN-TEx tissues with XCI ratios >= 0.7, we clearly see escape genes have balanced biallelic expression contributing to the potential for underestimating XCI ratios when including escape genes (Fig. 2e).

To assess XCI and escape more broadly, we capitalize on the tissue sampling structure of the Genotype-Tissue Expression (GTEx v8) dataset (Supp. Fig. 1). From an average of 56 +-23.5 (SD) well-powered heterozygous SNPs per sample (Supp. Fig. 2), we derive robust XCI ratio estimates for 4658 GTEx tissue samples spanning 49 different tissues (Supp. Fig. 3). Plotting the aggregated gene-level allelic expression ratios for all GTEx samples binned by tissue-level XCI ratio estimates reveals a heavy tail towards balanced allelic expression within more skewed tissues, indicative of novel escape genes (Fig. 2f).

### Escape genes exhibit consistent cross-tissue biallelic expression

Our method to quantitatively determine the tissue XCI ratio via aggregating signal across genes is especially well-suited to explore escape from XCI within the GTEx dataset (Fig. 2e). Our basic strategy for detecting escape genes is to calculate each gene’s consistency with the aggregate chromosomal XI ratio. Assessing all X-linked genes utilized in our GTEx XCI ratio estimates (Fig. 3a) and previously annotated constitutively escape genes^25^ results in a wide range of correlations between gene and tissue XCI ratios, exemplified by the genes SHROOM4 and TCEAL3 (Fig. 3b). As expected, the transcripts associated with XCI, namely, XIST and TSIX, show some of the highest correlations to the tissue XCI ratio (i.e., top 8.7%, Fig. 3b). Similarly, known escape genes exhibit some of the smallest correlations (Fig. 3b). Interestingly, several genes previously annotated as escape do exhibit rather strong correlations to the XCI ratio of tissues. We find that increased gene expression is linked to increased correlation to the tissue XCI ratio (Fig. 3c) suggesting that some gene variation with respect to the tissue XCI ratio is technical, reflecting read sampling at low expression. At matched expression levels, previously annotated escape genes have smaller tissue-gene XCI ratio correlations compared to all other genes (Fig. 3c), demonstrating that escape genes are less correlated to the tissue XCI ratio as expected by expression levels alone.

**Figure 3.**
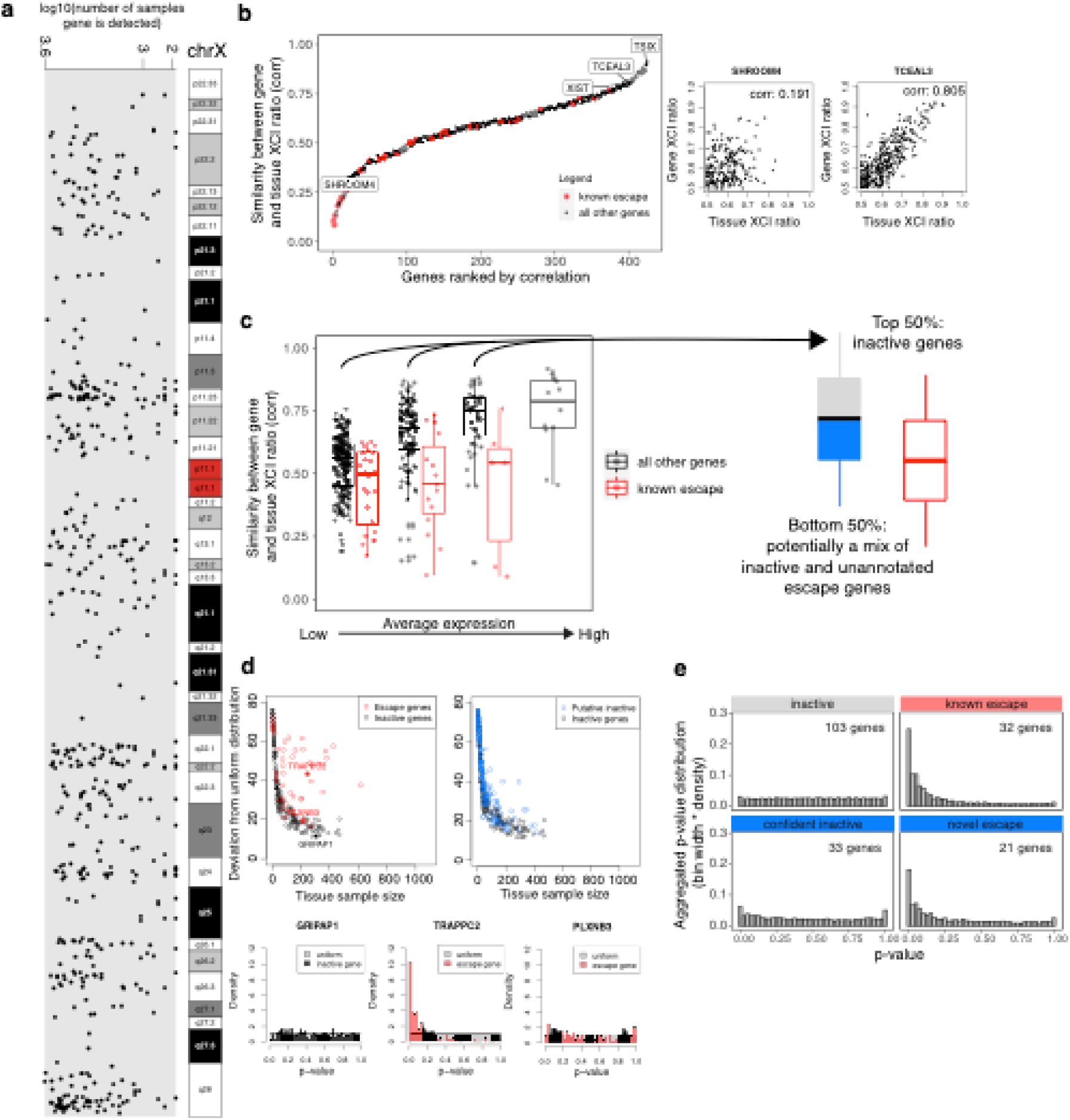
Genes that escape XCI exhibit balanced biallelic expression across XCI skewed tissues. **a**, The genomic location and number of GTEx samples each gene is detected for the 542 genes that pass our quality control filters. **b**, All 542 genes and 45 known escape genes ranked by the Pearson correlation coefficient for each gene’s allelic expression and the XCI ratio of the tissue for samples that detect that gene. **c**, Distributions of gene-tissue XCI ratio correlations for all 542 genes and 45 escape genes, binned by average expression. The range of average expression is binned into 4 equally spaced bins. For each expression bin, we labeled the top 50% of genes as inactive genes, genes that confidently follow the tissue XCI ratio, and the bottom 50% of genes as putative inactive genes, as they are potentially a mix of inactive and unannotated escape genes. **d**, The deviation from the uniform distribution is plotted for each gene’s p-value distribution, calculated from our one-sided test to detect balanced biallelic expression in tissues with XCI ratios >= 0.70 (see methods). The deviation of inactive genes was used as a threshold to identify robust escape genes and classify the putative inactive genes as either confident inactive or novel escape. **e**, The aggregated p-value distributions for inactive, known escape, and the putative inactive genes now classified as confident inactive and novel escape are plotted.

From our analysis in the EN-TEx dataset, we find escape from inactivation results in a bias for, rather than complete, balanced allelic expression (Fig. 2e), explaining how some escape genes retain significant correlations to tissue XCI ratios in the GTEx dataset. To comprehensively test the degree to which escape produces balanced allelic expression, we construct a one-sided test to detect whether a gene consistently trends towards balanced biallelic expression regardless of the XCI ratio of the tissue (see methods, Supp. Fig. 4). We restrict the tissue samples to have an estimated folded XCI ratio >= 0.7 to increase the power to detect balanced biallelic expression in opposition to the XCI ratio. Using this approach for the known escape genes results in the expected enrichment of significant p-values (Fig. 3d-e, Supp. Fig. 4). We apply stringent filters for calling escape status and select for genes that exhibit strong escape signal (Fig. 3 d). Some previously annotated escape genes that exhibit slight escape signal fail to pass our filters (PLXN3B, Fig. 3 d). This may reflect variation in escape across tissues.

Focusing on escape that generalizes across tissues, the genes previously unannotated for constitutive escape that display evidence for escape are ARHGAP4, ATRX, BTK, CASK, CLIC2, COX7B, CTPS2, CXorf36, EMD, F8, MECP2, MPP1, NLGN4X, PDZD4, PGK1, PLP1, RPL36A, SASH3, SEPT6, STARD8, VSIG4 (Fig. 3d-e, Supp. Fig. 4). Revisiting these genes within the literature, several have prior evidence for escape typically limited to a specific tissue: ATRX (cell lines^27^), BTK (PBMCs^28^), CLIC2 (escapes in skin^25^), COX7B (mouse embryonic stem cells^29^), CTPS2 (previous escape^30^), CXorf36 (escape in tumors^31^), NLGN4X (variable escape^25^), SEPT6 (B-cells^32^), VSIG4 (escape in mouse spleen^33^). Our results suggest these genes escape inactivation more broadly than previously reported. In addition, our analysis provides supporting evidence of escape for 32 previously annotated escape genes and supporting evidence of inactivation for 133 genes (Supp. Table 1).

### XCI is completed prior to germ layer specification

Having developed an approach to measure XCI ratios from unphased data, we turn to assessing the degree to which XCI ratios are shared across tissues within individuals. As an initial visualization of XCI ratios across tissues, we order all female GTEx donors by their average XCI ratio and plot the ratio for all tissues grouped by germ layer (Fig. 4a, Supp. Fig. 5). XCI ratios qualitatively appear consistent across all tissues and the three germ layers (Fig. 4a, Supp. Fig. 5). To test the degree of shared XCI ratios, we ask how well individual tissues predict highly skewed donor’s average XCI ratio across the remaining tissues (Supp. Fig. 6). If the XCI ratio is consistent across all tissues then every tissue is expected to perform well when predicting skewed donors. Across various thresholds for defining skewed donors, we see that performance is high and consistent across all tissues, suggesting XCI ratios are generally shared across all tissues for an individual (Supp. Fig. 6).

**Figure 4.**
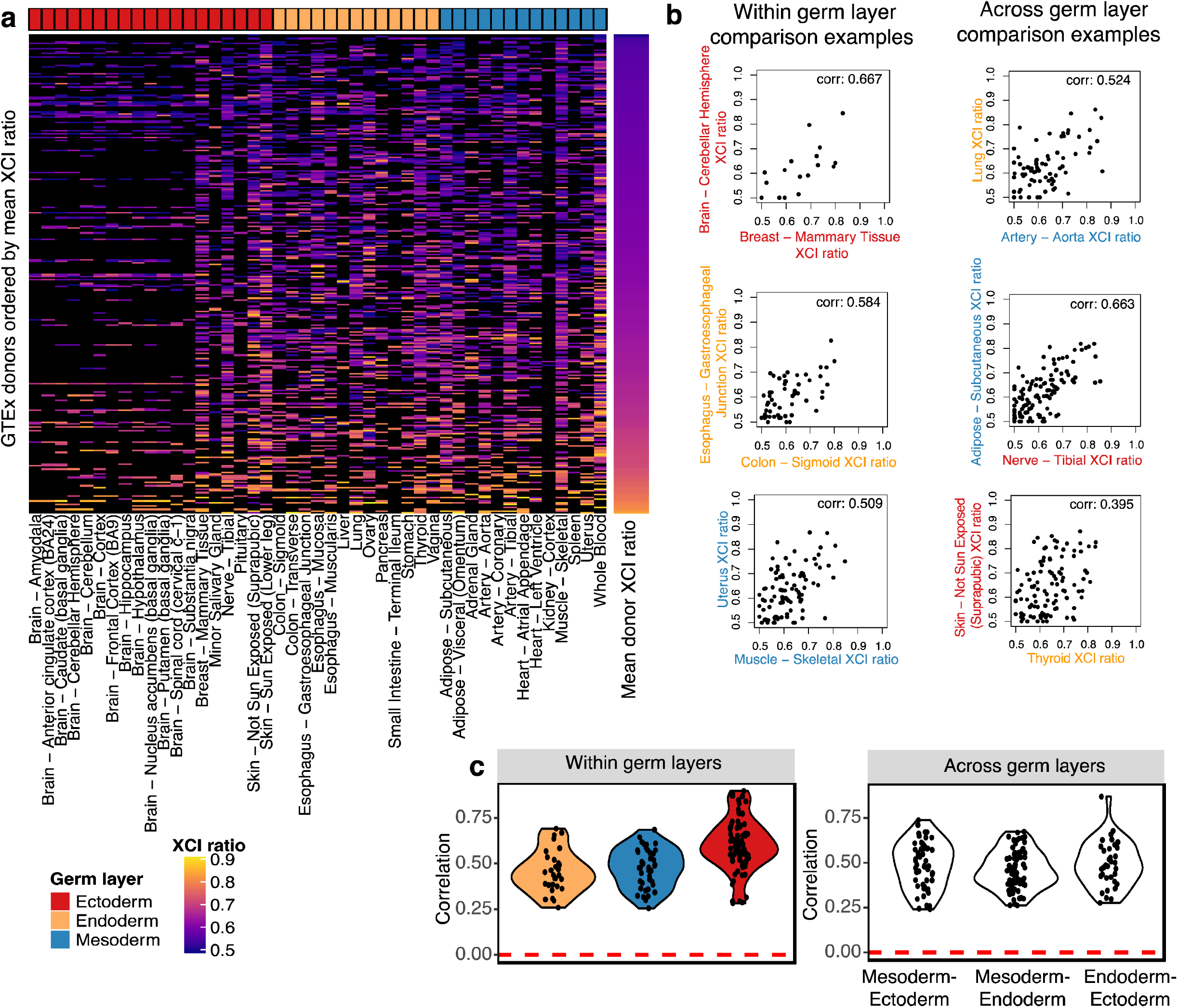
XCI ratios are shared across germ layer lineages. **a**, Heatmap of all estimated XCI ratios for the tissues of each donor, with donors ordered by their mean XCI ratio across tissues and tissues grouped by germ layer lineage. Black indicates no tissue donation for that donor-tissue pair. **b**, Examples of within and across germ layer lineage comparisons of XCI ratios. Each data point represents the estimated XCI ratios of the two indicated tissues for a single donor. **c**, All significant (FDR corrected p-value <= 0.5, permutation test n = 10000) Pearson correlation coefficients for within and across germ layer lineage comparisons.

Since folded ratios capture only the magnitude of the tissue XCI ratio, two tissues with XCI ratios in opposite parental directions might appear similar. To investigate whether the parental direction of XCI differs for the same individual between two tissues, we turn to shared SNP-level allelic ratios across tissues for individual donors. If an expressed heterozygous SNP is captured for two different tissues of an individual, the reference allele is on the same haplotype and maintains directional allelic information. Thus, calculating the correlation of reference SNP allelic ratios for shared SNPs between two tissues can reveal whether those tissues share the same XCI direction (Supp. Fig.7). For donors with at least 10 tissues, the average percent of tissues exhibiting switched XCI is 8.3%, suggesting switched XCI occurs infrequently. Interestingly, XCI switching is concentrated in a few tissues with 12 out of 49 being significantly enriched for instances of switched XCI (Supp. Fig. 7). This holds true for the phased EN-TEx data as well, where only the liver, ovary, and omental fat pad change parental XCI ratio directions (Supp. Fig. 7). One plausible explanation for this effect is that the XCI ratio for some tissues is subject to additional sources of variability compared to other tissues, such as lineage specific sampling (See section ‘Cell population estimates at the time of tissue-specific lineage commitment’).

The broad sampling of different tissues present within the GTEx dataset enables a well-powered investigation of shared XCI ratios across tissues. Importantly, stratifying these tissue comparisons by germ layer lineage relationships should resolve the temporal ordering of XCI and germ layer specification within the human embryo. If XCI occurs before germ layer specification, tissue XCI ratios are expected to positively covary across tissues from different germ layer lineages (Fig.1a). In contrast, if XCI occurs after germ layer specification, the XCI ratio of each germ layer is set independently and there is little expected covariance in XCI ratios for tissues from different germ layers (Fig.1b). We compute correlations of the XCI ratio for combinations of tissues derived from either the same or different germ layers, exemplified in Figure 4 panel b. Tissues sharing the same germ layer lineage produce strictly positive significant correlation values ranging from 0.25 to 0.90 (Fig. 4c1), demonstrating XCI ratios are shared within individual germ layer lineages. Strikingly, significant positive ratio correlations for tissues derived from different germ layers are on the same order as the within germ layer comparisons, ranging from 0.24 to 0.87 (Fig. 4c2, Supp. Fig. 6). The fact tissues derived from different germ layers covary for their XCI ratio strongly suggests XCI is completed prior to germ layer specification and the initial embryonic XCI ratio is propagated through all germ layer lineages.

### Cell population estimate at the time of XCI

The fact XCI ratios are shared across tissues suggests the initial embryonic XCI ratio determined at the time of inactivation is propagated through development. Population level variance in adult XCI ratios thus reflects the sample distribution during XCI, which depends on the number of cells present during inactivation. We derive estimates for the number of cells present at the time of inactivation by modeling XCI ratio variance from tissue-specific ratio distributions across donors (Fig. 5a, Supp. Fig. 8). Using a maximum likelihood approach, we fit estimated models to the tails of the empirical XCI ratio distributions to account for the uncertain unfolded XCI ratio estimates between 0.4 and 0.6 (Fig. 5a, see methods). The cell number estimates derived from all well-powered tissues range from 6 to 16 cells (Fig. 5b), i.e., approximately within a single cell division, demonstrating a striking degree of similarity in population level XCI ratio variance across the assessed tissues. By our estimates, human XCI is completed at a time in the embryo when between 6 and 16 cells are fated to become all embryonic tissues.

**Figure 5.**
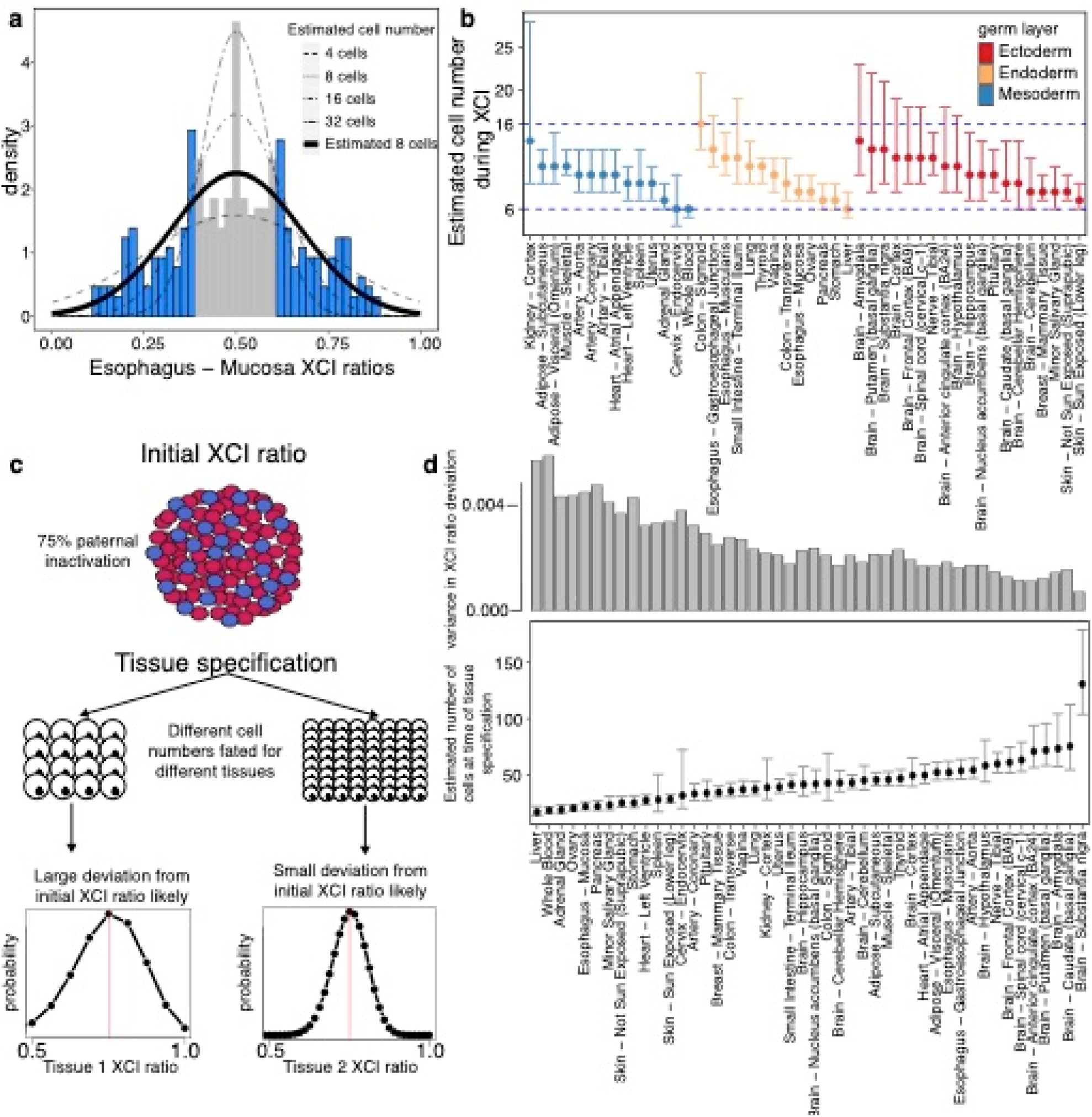
XCI and tissue lineage specification can be timed to a pool of cells by exploiting observed variability. **a**, Example tissue demonstrating the model for estimating cell numbers at the time of XCI using the population-level variance in XCI ratios. We fit normal distributions, as a continuous approximation of the underlying binomial distribution of XCI ratios, to the tails of tissue-specific XCI ratio distributions (shaded in blue), which accounts for the uncertain 0.40-0.60 unfolded XCI ratio estimates (shaded in grey). **b**, The resulting estimated cell numbers present during XCI derived from the XCI ratio variance of all tissues with at least 10 donors. Error bars are 95% confidence intervals and tissues are grouped by germ layer lineage. Cell number estimates range from 6-16 cells across all tissues. **c**, Schematic for our model of tissue lineage specification and the implications for tissue-specific XCI ratios. The XCI ratio of a tissue is dependent on the prior XCI ratio of the embryo and the number of cells selected for that tissue lineage. These two features define the binomial distribution for that tissue’s XCI ratio. **d**, Estimated number of cells selected for individual tissue lineage specification of 46 different tissues. Error bars represent 95% confidence intervals. The top bar graph plots the variance in the distribution of tissue XCI ratio deviation from the average XCI ratio of each donor for that tissue. Our model follows the logic that lineage specification of a small pool of cells increases variation in XCI ratios, resulting in increased likelihood a tissue will deviate from the average donor XCI ratio.

### Cell population estimates at the time of tissue-specific lineage commitment

Tissue-specific lineage commitment can be modeled as a random sampling event from a pool of unspecified progenitor cells. In the context of XCI, the XCI ratio of the newly specified tissue is dependent on the prior XCI ratio of the progenitor pool and the number of cells fated for that tissue and can be modeled as a binomial sampling event (Fig. 5c). As such, the GTEx dataset offers a unique opportunity to capture this tissue-specific XCI variance and model the number of cells present at the time of tissue-specific lineage commitment across a broad range of human tissues.

To capture the tissue-specific variance in XCI as it relates to the prior embryonic XCI ratio, we model the deviation of tissue-specific XCI ratios from the average donor XCI ratios for all donors of a given tissue (see Methods, Fig. 5d, 46 well-powered tissues). Our model follows the logic that tissues with large variation in their deviation from average donor XCI ratios are derived from a smaller pool of cells, a consequence of increased variability due to small sample size effects. Intriguingly, we find a strong negative relationship between our estimated cell numbers at the time of tissue-specific lineage commitment and the probability of a tissue switching the direction of XCI in comparison to other tissues of an individual (Supp. Fig. 7). This is in line with the model that tissues derived from smaller stem cell pools are subject to increased XCI ratio variability and thus more likely to produce a sample of cells with their XCI ratio in the opposite direction from the prior embryonic XCI ratio.

## Discussion

In this work, we exploited the random, permanent, and developmentally early nature of XCI to investigate characteristics of early lineage specification events during human development. By analyzing variance in XCI ratios across tissues and individuals, we showed human XCI is completed before tissue specification and the stochastically determined XCI ratio set during embryogenesis is a shared feature across all tissue lineages. We estimate between 6-16 cells are fated for all embryonic tissues at the time of XCI completion. Additionally, we provide estimates of the number of cells present during tissue-specific lineage specification for 46 different tissues. To conduct this analysis, we developed a method to estimate the ratio of XCI using unphased allele-specific expression, a highly scalable approach applicable to any bulk RNA-sequencing sample.

As our approach to estimate XCI ratios is independent of the underlying genes used to assess allele-specific expression, it can be applied across systems with variable expression profiles, such as tissues, for comparative XCI analysis. As such, our method is well-suited for taking advantage of the extensive cross-tissue data of the GTEx dataset and for extracting the whole-body lineage relationships present therein. We show that XCI ratios are shared across all tissues for an individual, providing insight into the observed variance of XCI ratios in normal female populations, an area of ongoing debate^10,34,35,36^. Our results indicate that the initial embryonic XCI ratio is propagated through development and is a shared feature across all tissue lineages. This demonstrates that the stochasticity of the initial choice for inactivation within the embryo has a measurable impact on XCI ratios in adult females. Additionally, our results determining the number of cells fated for tissue-specific lineage commitment revealed a broad range of cell numbers, identifying an additional layer of variation where developmental stochasticity can have a phenotypic impact. Tissues derived from smaller stem cell pools may be more susceptible to stochastic developmental events, such as deleterious *de novo* mutations, a possibility strongly suggested by our finding that tissues with small estimated stem cell pools are more likely to switch the direction of XCI within an individual.

Our work is part of a broader history of using X-linked mosaicism as a tool for studying lineage relationships, with studies ranging from investigations of early lineage events in mice^6^ to ascertaining tumor clonality^37^. Typically, these approaches will capitalize on a single locus of the X-chromosome to determine XCI status^38^. When using allele-specific expression, the use of a fully phased genome would similarly allow any single heterozygous X-linked loci to mark the expressed X, though generating a phased genome is generally cost prohibitive. Importantly, the variability in expression across systems and the fidelity of inactivation for individual genes (escape from XCI) means that robust cross-tissue evaluation of XCI using single X-linked loci has been challenging to conduct. One of our methodological contributions is to demonstrate the allelic expression imbalance generated via XCI can be aggregated to provide near-perfect estimates of XCI ratios, even in the absence of phased information. Indeed, while none of our estimated XCI ratios are phased at the SNP-level, the observed error in our estimates by aggregating allelic expression correctly across SNPs is lower than any single locus can provide.

While GTEx represents the premier dataset for human cross-tissue functional genomics, more data is always helpful. As our approach for estimating XCI ratios is applicable to any bulk RNA-sequencing data, we envision this work providing an informative control for any future functional genomic investigations involving the X-chromosome.

## Methods

### GTEx and EN-TEx data

Fastq files for all female donors from the GTEx project v7 release^20^ were obtained from dbGaP accession number phs000424.vN.pN. BAM files for additional female samples from the v8 release were obtained from the associated AnVIL repository (gtexportal.org/home/protectedDataAcccess). Phased expression data from the EN-TEx project^23,24^ were obtained in collaboration with the ENCODE consortium. EN-TEx data is available on the online portal^23^.

### RNA-seq alignment and SNP identification

For aligning RNA-sequencing data, the GRCh38.p7 human reference genome using GENCODE v.25 annotations was generated with STAR v2.4.2a^39^ and data was aligned with STAR v2.4.2a or STAR v2.5.2b. STAR was run using default parameters with per sample 2-pass mapping. BAM files for the additional GTEx v8 samples (originally aligned to GRCh38.p10 with GENCODE v.26 annotations) were sorted using samtools v1.9^40^ and converted to fastq files using bedtools v.2.26.0^41^. For each sample, alignment to the X-chromosome was extracted using samtools and passed to GATK^42,43^ for SNP identification. Using GATK v.4.1.3.0 and following the best practices workflow for RNAseq short variant discovery (<u>GATK best practices</u>), we used the following pipeline of GATK tools using default parameters unless otherwise stated: AddorReplaceReadGroups -> MarkDuplicates -> SplitNCigarReads -> HaplotypeCaller (-stand-call-conf 0.0) -> SelectVariants (-select-type SNP) -> VariantFiltration. The following filters were used in VariantFiltration to set flags for downstream filtering: QD < 2.0, QUAL < 30.0, SOR > 3.0, FS > 60.0, MQ < 40.0, MQRankSum < -12.5, and ReadPosRankSum < -8.0. These filters were determined from GATK recommendations and empirical evaluation of the identified SNPs’ metrics.

### SNP quality control

SNPs identified through GATK were further filtered by various metrics to increase confidence in SNPs identified from RNA-sequencing data and ensure well-powered SNPs for allele-specific expression analysis. The resulting .vcf files from GATK were filtered to only contain SNPs present within dbSNP^44^. The remaining SNPs were filtered to be heterozygous with 2 identified alleles and at least 10 reads mapped to each allele for a minimum threshold of 20 reads per SNP. Additionally, SNPs were required to pass the SOR, FS, and ReadPosRankSum filters set in the GATK pipeline. Only SNPs located within annotated genes (excluding the PAR regions of the X-chromosome) were considered and in the case of multiple identified SNPs in the same gene for a sample, the SNP with the highest total read count was taken as the max-powered representative for that gene. SNPs with a total read count above 3000 were excluded as they demonstrated a uniform distribution of allelic expression (Supp. Fig. 2).

### Gene filtering (reference bias and XCI escape)

From the observation of a heavy tail towards allelic expression in the reference direction across all called SNPs in the GTEx dataset (Supp. Fig. 2), we compiled gene specific distributions of allelic expression to determine if a select few genes/SNPs were at fault. The majority of genes demonstrated distributions of relative allelic expression centered around 0.5 with several considerable exceptions, some genes exhibited bimodal or extremely biased distributions (Supp. Fig. 2). We excluded genes that failed the dip test for bimodality as well as the top and bottom 5% of genes ranked by the deviation of their mean reference expression ratio from 0.5 (Supp. Fig. 2). Additionally, we excluded genes previously annotated to constitutively escape XCI^25^. In total, we end up with well-powered SNPs from 542 genes along the X-chromosome for modeling XCI ratios.

### Folded normal model for estimating XCI ratios

We aggregate the allelic expression imbalance of the X-chromosome over both alleles by folding the reference allelic expression ratios about 0.5 (Fig 2a-b). To obtain our XCI ratio estimates we fit a folded normal distribution to the folded reference allelic expression ratios of each sample, using the maximum log likelihood estimate as the estimated XCI ratio. Theoretically, the captured bulk allelic expression for a heterozygous X-linked SNP follows a binomial distribution characterized by the read depth of the SNP and the XCI ratio of the sample. Without phasing information, the allelic expression of heterozygous X-linked SNPs can be characterized by the folded-binomial model^21,22^. Since SNPs vary in read depth and various biological factors (e.g. eQTLs) are not accounted for in the binomial model, we take the folded normal model as a continuous approximation. We require samples to have XCI ratio estimates derived from at least 10 filtered SNPs for downstream analysis, resulting in 4659 samples with a mean of 56 well-powered SNPs per sample (Supp. Fig. 3). Additionally, we calculate 95% confidence intervals (CI) for each XCI ratio estimate via a nonparametric bootstrap percentile approach (n = 200), excluding XCI ratio estimates with a CI width >= .15 from downstream analysis. For donors with multiple samples for the same tissue, we average the XCI ratio estimates together, duplicated tissue samples have minor differences in estimated XCI ratios (mean difference in XCI ratios for duplicate tissue samples: 0.018 +-0.023 SD).

### Gene-tissue XCI ratio correlations

To test an individual gene’s propensity to follow the aggregate chromosomal XCI ratio, we calculate Pearson correlations between a gene’s reference allelic expression ratio and the estimated XCI ratio (leaving out that gene for all samples in which the gene is detected). We calculate these correlations for each of the 542 filtered genes described above and for 45 previously annotated constitutively escape genes detected in our dataset. We only consider genes detected in at least 30 samples and with an FDR corrected (Benjamini-Hochberg) correlation p-value <= .05 determined by a permutation test (n = 10000) for further investigation of escape status, resulting in 380 putative inactive genes and the 45 previously annotated escape genes.

### Testing for escape from XCI

To detect escape genes, it is necessary to compare against genes that undergo complete inactivation and do not escape. After they are stratified by mean expression, we reason the genes most likely to undergo complete inactivation are genes with high gene-tissue XCI ratio correlations within each expression bin (Fig. 3c). Accordingly, we take the top 50% of putative inactive genes within each bin to define the null distribution of allelic expression under the hypothesis of complete inactivation (191 genes). The remaining 189 putative inactive genes and the 45 known escape genes composes our test set. We reason a gene that escapes XCI will be biased for balanced biallelic expression regardless of the XCI ratio of the tissue. Using only tissues with an estimated XCI ratio >= 0.70, we compute the deviation from 0.5 (balanced allelic expression) for all genes and calculate the empirical p-value as follows: sum (deviation (null inactive genes) <= deviation (test gene)) / number of genes (Supp. Fig. 4). We only consider p-values derived from samples with at least 20 null inactive genes detected.

Under the null hypothesis of inactivation, genes with strong deviation from a uniform p-value distribution are strong candidates for escape from inactivation. We first fit a slightly offset negative exponential function to the uniform deviations of the null inactive genes to define a threshold for escape while controlling for sample size (Supp. Fig. 4). Additionally, we only consider genes with at least 50 computed p-values. An enrichment of low p-values from this test is indicative of balanced biallelic expression regardless of the XCI ratio of the tissue. It follows that an enrichment of high p-values from this test is indicative of preferential mono-allelic expression regardless of the XCI ratio of the tissue.

### Tissue XCI ratio predicting donor XCI ratio

For the donors that contribute to a given tissue, we calculate the mean XCI ratio across all other tissues for each donor. We define skewed donors as having a mean XCI ratio greater than or equal to various thresholds (0.65, 0.7, 0.75, 0.8). We calculate the AUROC of a given tissue’s XCI ratio predicting skewed donors via the Mann-Whitney U test statistic.

### Cross-tissue XCI ratio correlations

For all pairwise combinations of the 49 tissues present within the GTEx dataset, we take the subset of donors that contribute both tissues for a given comparison and calculate the Pearson correlation for the folded XCI ratio of the tissues. Figure 4c1-c2 depicts only the correlation values derived with a sample size of at least 20 donors and an FDR corrected (Benjamini-Hochberg) p-value <= .05 derived from a permutation test (n = 10000). Supplemental Figure 6 depicts all computed correlations regardless of sample size or p-value.

### Inference on direction of XCI ratios

To infer the direction of XCI ratios from unphased data, we look at allelic expression of heterozygous SNPs captured in multiple tissues for an individual donor. The reference allele of a heterozygous SNP captured in two different tissues of a single donor represents the same parental X-allele in both tissues. If the direction of XCI is the same for both tissues, the heterozygous SNP is expected to exhibit the same degree of reference allelic expression across the two tissues (positive correlation). If the direction of XCI is different, reference allelic expression will be inverted for one of the tissues resulting in a negative correlation. For each donor, for all pairwise combinations of their donated tissues with XCI ratios >= 0.6, we calculate Pearson correlations for unfolded reference allelic expression ratios using only SNPs detected in both tissues (Supp. Fig. 7). We only use SNPs that are within the previously filtered 542 genes described above and only consider correlations derived from tissue comparisons with at least 30 shared SNPs. Using positive or negative correlations as a readout for switched XCI direction between tissues, we perform Fisher’s exact test with a Benjamini-Hochberg correction to identify any tissue significantly enriched for switching XCI directions.

### Evaluating XCI cell number estimates

XCI is a binomial sampling event defined by the number of cells present during inactivation and the equal probability of inactivation between the alleles (N = # of cells, p = 0.5). As such, the variance in XCI ratios within a population is directly linked to the number of cells present during XCI. We derive estimates for the number of cells present during XCI by fitting a normal model to tissue-specific XCI skew distributions as a smoothened estimate for the underlying binomial distribution. We take the theoretical variance from the binomial model as the variance for the normal approximation.

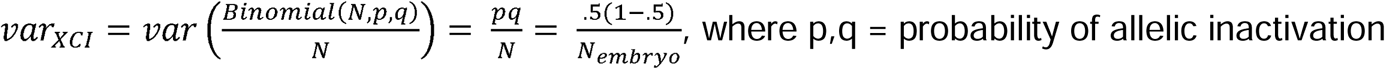

For a range of cell numbers (N = 2:50), we select the normal model with minimum error between its CDF and the empirical XCI ratio CDF of a given tissue for the tails of the distribution (XCI ratio <= 0.4 and XCI ratio >= 0.6, Supp. Fig. 8). This accounts for the uncertain folded 0.5 – 0.6 XCI ratios estimates in the unfolded space. We calculate 95% CIs for each estimated cell number via a nonparametric bootstrap percentile approach (n = 2000). We only consider cell number estimates from tissues with at least 10 donors.

### Evaluating tissue-specific lineage cell number estimates

We model tissue-specific lineage specification as a cell sampling event from a large pool of cells. As such, the XCI ratio of a tissue will follow a binomial model defined by the number of cells fated for that tissue and the XCI ratio of the embryo (Fig. 5c).

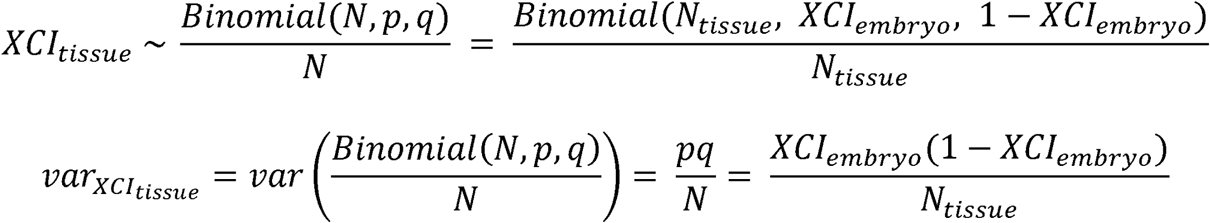

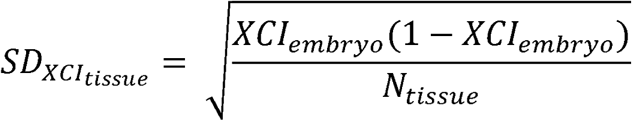

For a given tissue, across donors with variable XCI ratios (*XCI*_*embryo*_) the variation in the tissue XCI ratio is defined by the constant *N*_*tissue*_, the number of cells fated for that tissue. To estimate this constant, we calculate z-scores for each tissue-donor pair of a given tissue using the mean XCI ratio of all other tissues for each donor as an approximation for *XCI*_*embryo*_.

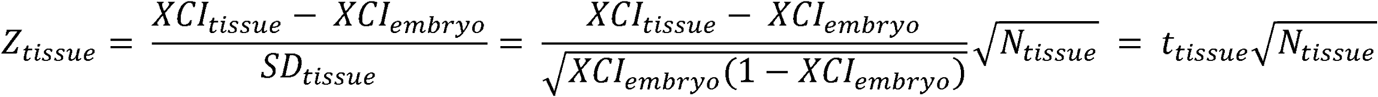

As the standard deviation of a distribution of z-scores is 1, we solve for *N*_*tissue*_:

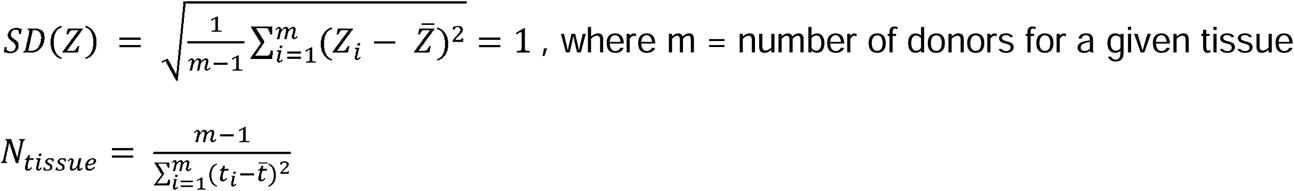

We calculate 95% CIs for each *N*_*tissue*_ via a nonparametric bootstrap percentile approach (n = 2000) using the *t*_*tissue*_ distribution. We require a tissue to have at least 10 donors in order to calculate *N*_*tissue*_.

### Data analysis and visualization

All analysis was conducted in R version 4.0.5^45^. Graphs were generated using the ggplot2^46^, ComplexHeatmap^47^, karyoploteR^48^, and base R packages.

### Data and code availability

The code for all analysis is available on Github github.com/JonathanMWerner/human_cross_tissue_XCI. Processed data is available on the FTP site http://labshare.cshl.edu/shares/gillislab/people/werner/xskew2021_preprint/data.

## Supporting information

Supplemental Figures with legends

## Acknowledgements

JMW was supported by NSF Award No. DGE-1938105. JG, SB and JH were supported by NIH Grants R01MH113005 and R01LM012736.

This material is based upon work supported by the National Science Foundation Graduate Research Fellowship Program under Grant No. DGE-1938105. Any opinions, findings, and conclusions or recommendations expressed in this material are those of the author(s) and do not necessarily reflect the views of the National Science Foundation.

We thank all members of the Gillis lab and particularly Manthan Shah for assisting in some of the initial downloading.

## Author Contributions

JG conceived the project. JMW and JG designed the experiments and wrote the manuscript. JMW performed the experiments. SB and JH assisted with data management and other initial data curation.

