## Supplemental Figures with legends for "Cross-tissue analysis of allelic X-chromosome inactivation ratios resolves features of human development"

**Author information**

1 The Stanley Institute for Cognitive Genomics, Cold Spring Harbor Laboratory, Cold Spring Harbor, NY, 11724, USA

2 Garvan Institute of Medical Research, Darlinghurst, Sydney, Australia

**Supplemental information**

**Supplemental Table 1**

Escape annotations

Assigned labels of inactive, known escape, confident inactive, or novel escape for the 189 genes powered enough to investigate cross-tissue XCI escape.

**Supplemental Figure 1**

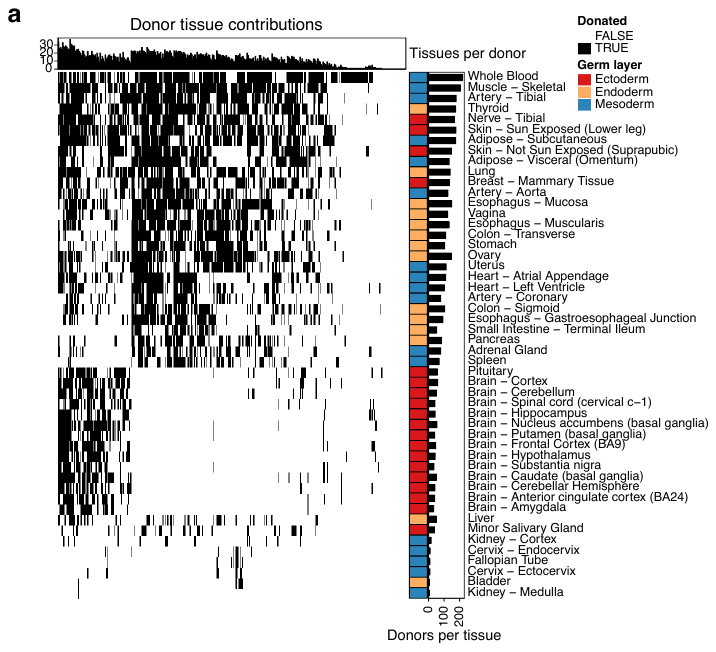

GTEx represents tissues across the three germ layer lineages for hundreds of female donors

**a**, Binary heatmap of female donor tissue contributions in the GTEx dataset for samples that pass our quality control filters. Data from cell lines was excluded in the final analysis.

**Supplemental Figure 2**

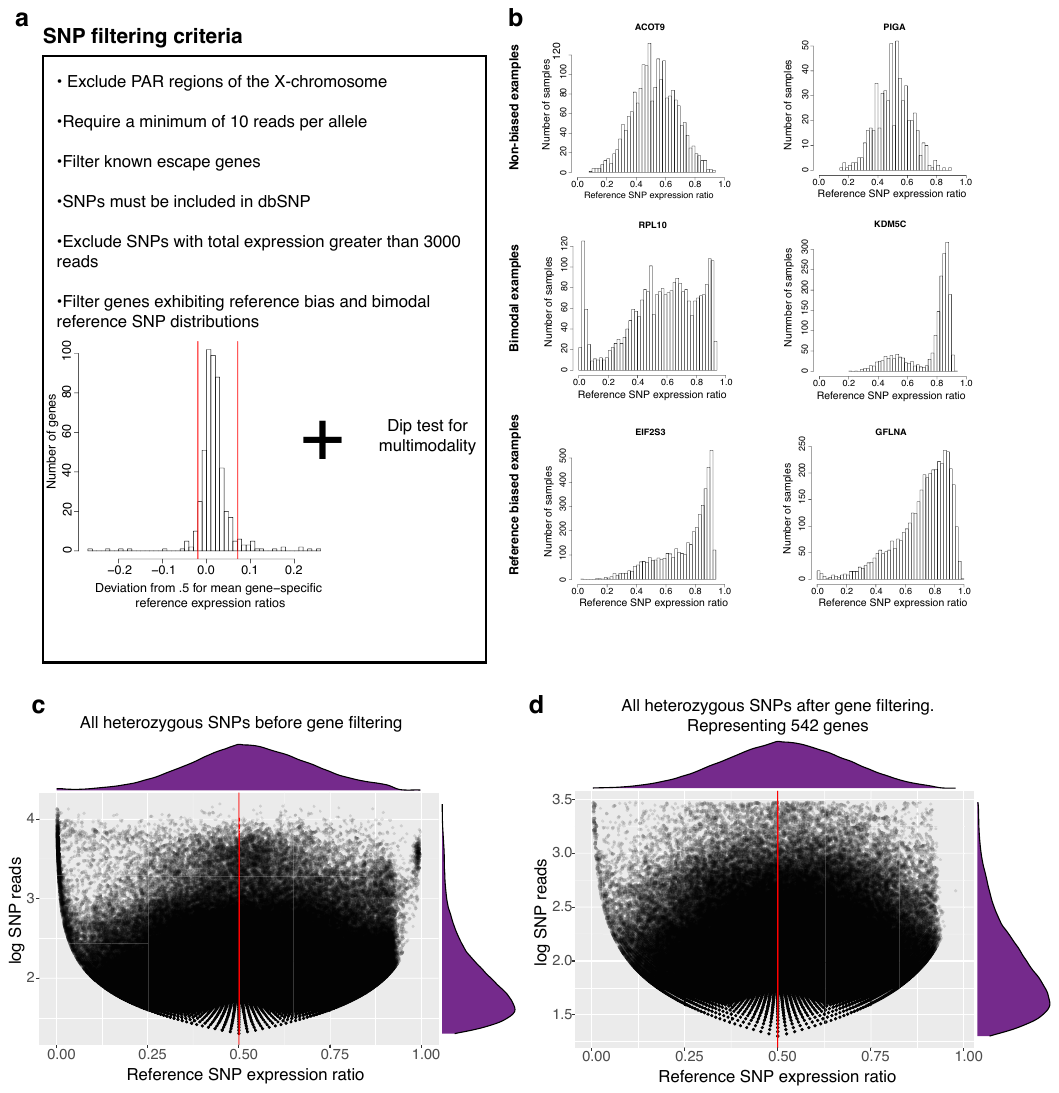

Rigorous quality control filters identify well-powered heterozygous SNPs from RNA-sequencing data

**a**, Bulleted list of filters for individual SNPs and genes. Includes histogram of the deviation from 0.5 for the mean reference expression ratios of individual genes, indicative of consistent gene-specific reference or alternate allele bias. Red horizontal lines indicate thresholds for excluding biased genes. **b**, Examples of reference allelic expression ratio distributions for individual genes, highlighting examples of genes with bimodal or reference biased distributions. **c**, Scatter plot with marginal density plots of reference allelic expression ratios and total reads for all identified heterozygous SNPs with at least 10 reads per allele prior to gene filtering. There is a heavy tail in the reference direction indicative of reference bias. **d**, Same plot as in c but after gene filtering.

**Supplemental Figure 3**

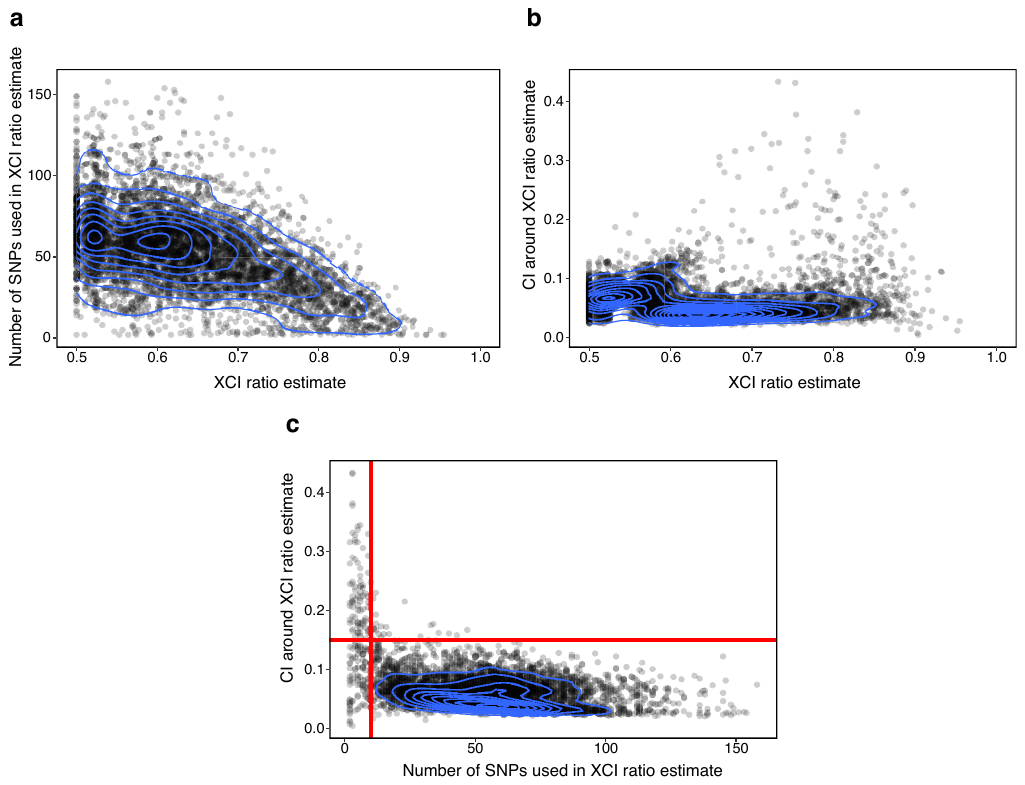

Filtering tissue XCI ratio estimates

**a**, Scatter plot with 2d density overlay of all XCI ratio estimates for 5046 GTEx samples and the number of filtered heterozygous SNPs used to estimate the sample XCI ratio. **b**, Scatter plot with 2d density overlay of all XCI ratio estimates for 5046 GTEx samples and the width of the 95% confidence interval around the XCI ratio estimate (bootstrap sampling, n = 200). **c**, Scatter plot with 2d density overlay of the number of filtered heterozygous SNPs used to estimate the sample XCI ratio and the width of the 95% confidence interval around the XCI ratio estimate. Red lines indicate thresholds for XCI ratio estimate filtering, requiring >= 10 heterozygous SNPs and a CI width < 0.15.

**Supplemental Figure 4**

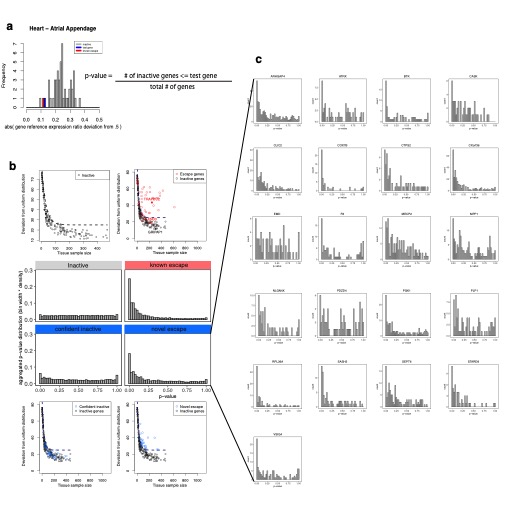

Genes that escape XCI are identifiable via balanced allelic expression in skewed XCI tissues

**a**, Histogram of gene reference allelic expression ratio deviations from 0.5 for a sample with an estimated XCI ratio >= 0.70. An example known escape gene in the sample is colored red and an example putative inactive gene is colored blue. Includes the formula for calculating the empirical p-value for a gene under the null hypothesis of inactivation for a single tissue sample. **b**, Central histograms are the same plots as in Figure 3e. Surrounding scatter plots compare the number of tissue samples a gene was detected, i.e., the number of p-values that make up each gene’s p-value distribution under our test for escape, compared to the deviation from the uniform distribution of each gene’s p-value distribution. A slightly offset negative exponential function was fit to the inactive to define a threshold for calling escape from XCI while controlling for sample size **c**, p-value distributions for the 21 genes that we classify as novel escape genes, as they exhibit significant signal under our test for escape.

**Supplemental Figure 5**

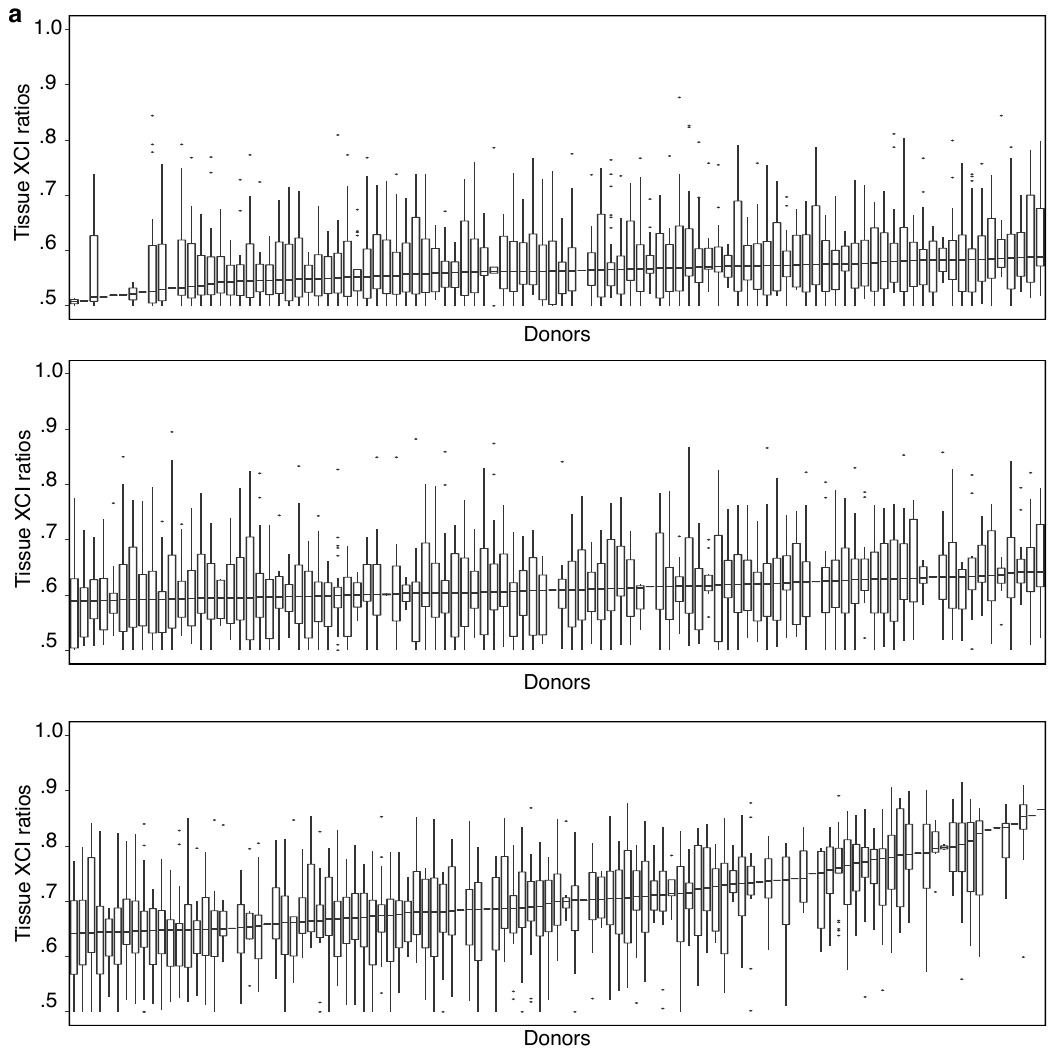

XCI ratios are shared across tissues

**a**, Boxplots of XCI ratios for all tissues donated by each individual donor (n = 311). Donors are ordered by their median tissue XCI ratio.

**Supplemental Figure 6**

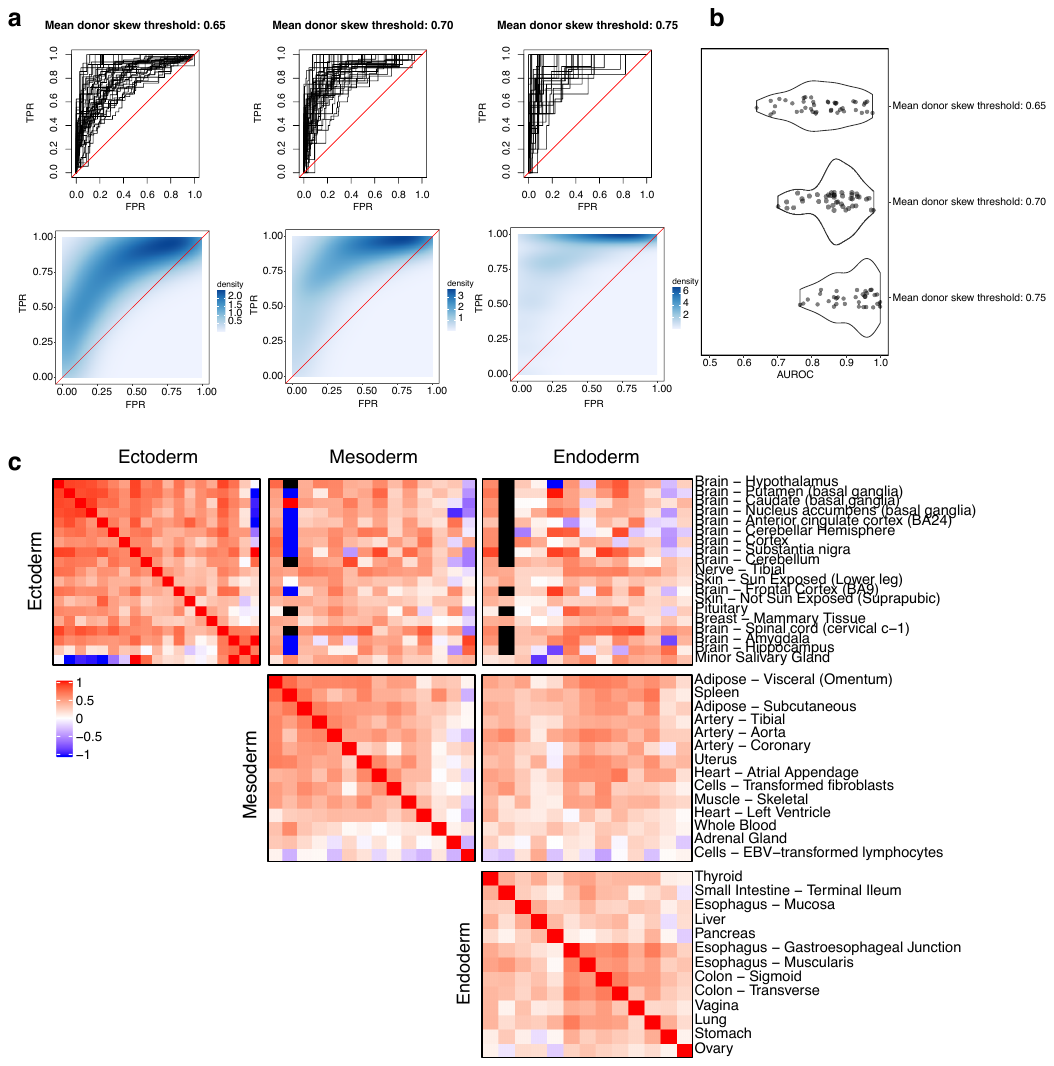

All tissues can strongly predict skewed donors and are correlated in XCI ratios

**a**, ROC curves for individual tissue XCI ratios predicting skewed donors at various thresholds for classifying skewed donors (top row). 2d density estimations across all tissue ROC curves (bottom row). **b**, AUROC distributions at each skewed donor threshold. **c**, All pairwise tissue-tissue XCI ratio correlations regardless of sample size or significance, grouped by germ layer lineage. The global trend is a positive correlation.

**Supplemental Figure 7**

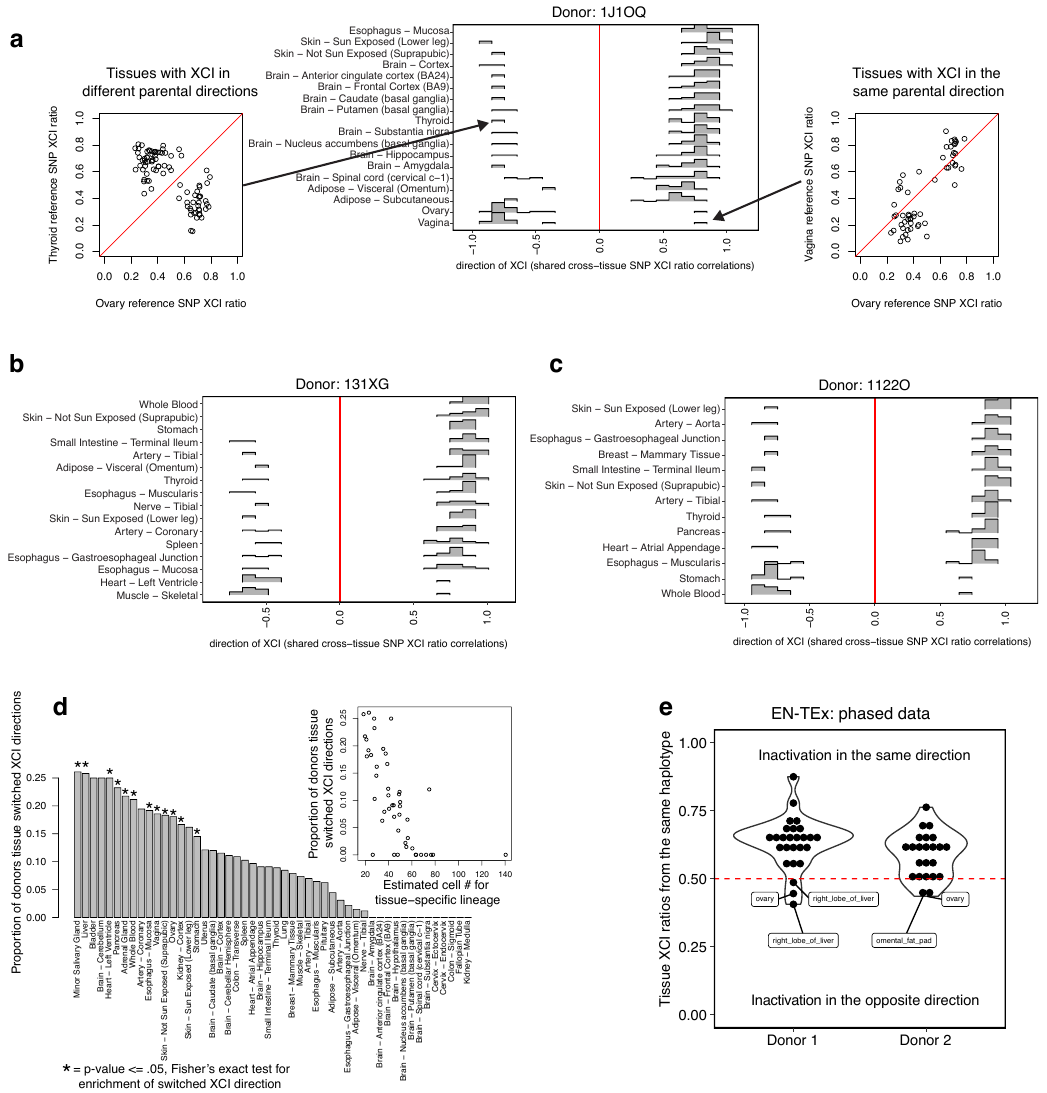

Tissues share the same allelic direction of XCI within an individual

**a**, Leftmost scatter plot contains reference SNP allelic expression for identical SNPs captured in two different tissues for a single donor. A negative linear correlation (Pearson) indicates the direction of XCI is different between the two tissues. Rightmost scatter plot contains the same data for two different tissues with a positive linear correlation, indicative of a shared XCI direction. Central histograms are the Pearson correlation distributions for all pairwise comparisons of tissues for a single donor. **b**, **c**, Same plot as the central histograms in a, but for 2 additional donors with more than 10 tissues as examples. **d**, Bar plot indicating the proportion of donors where the specified tissue switched directions compared to other tissues. Asterisks indicate significance from Fisher’s Exact test (FDR corrected p-value <= .05), identifying tissues enriched for switching XCI directions. Inset scatter plot compares the estimated cell number for tissue specific lineage specification from Figure 5d to the proportion of switched XCI for each tissue. **e**, Violin plots of median phased gene XCI ratios from the same haplotype for each tissue of both EN-TEx donors.

**Supplemental Figure 8**

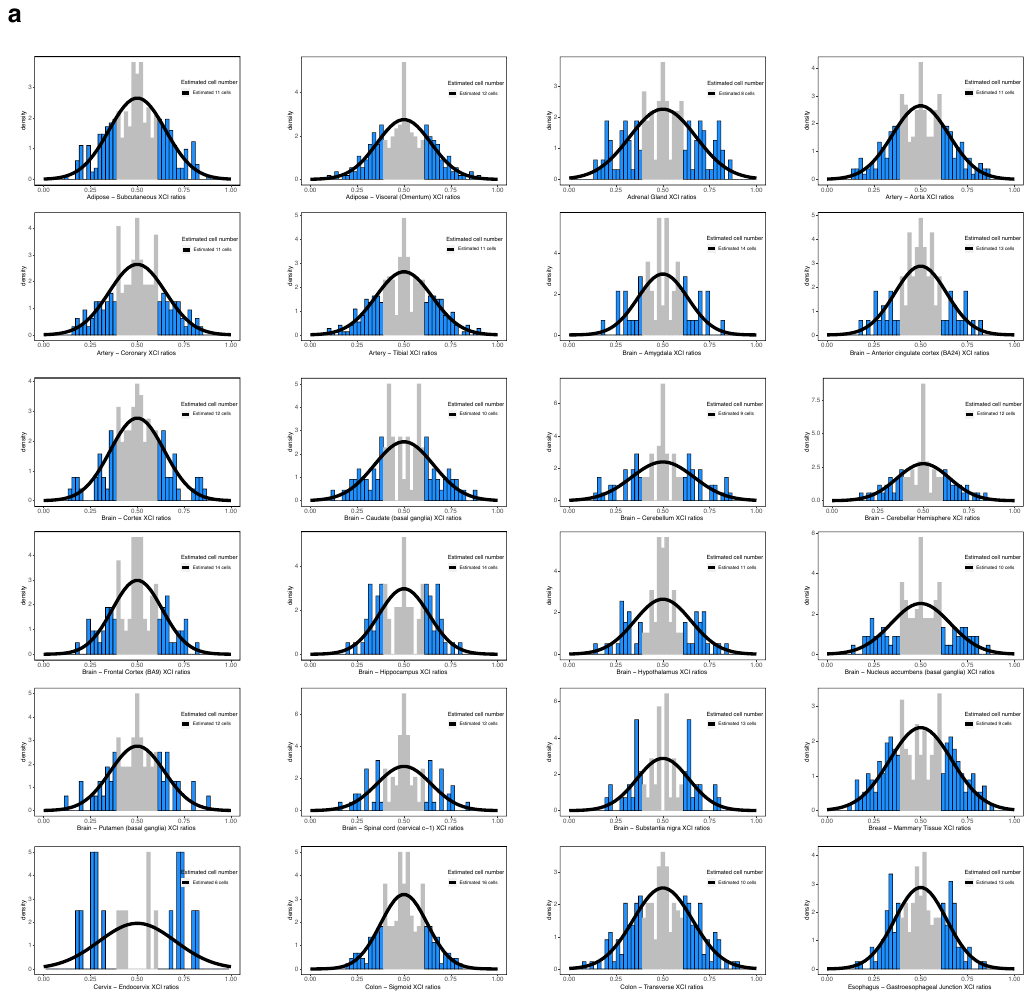

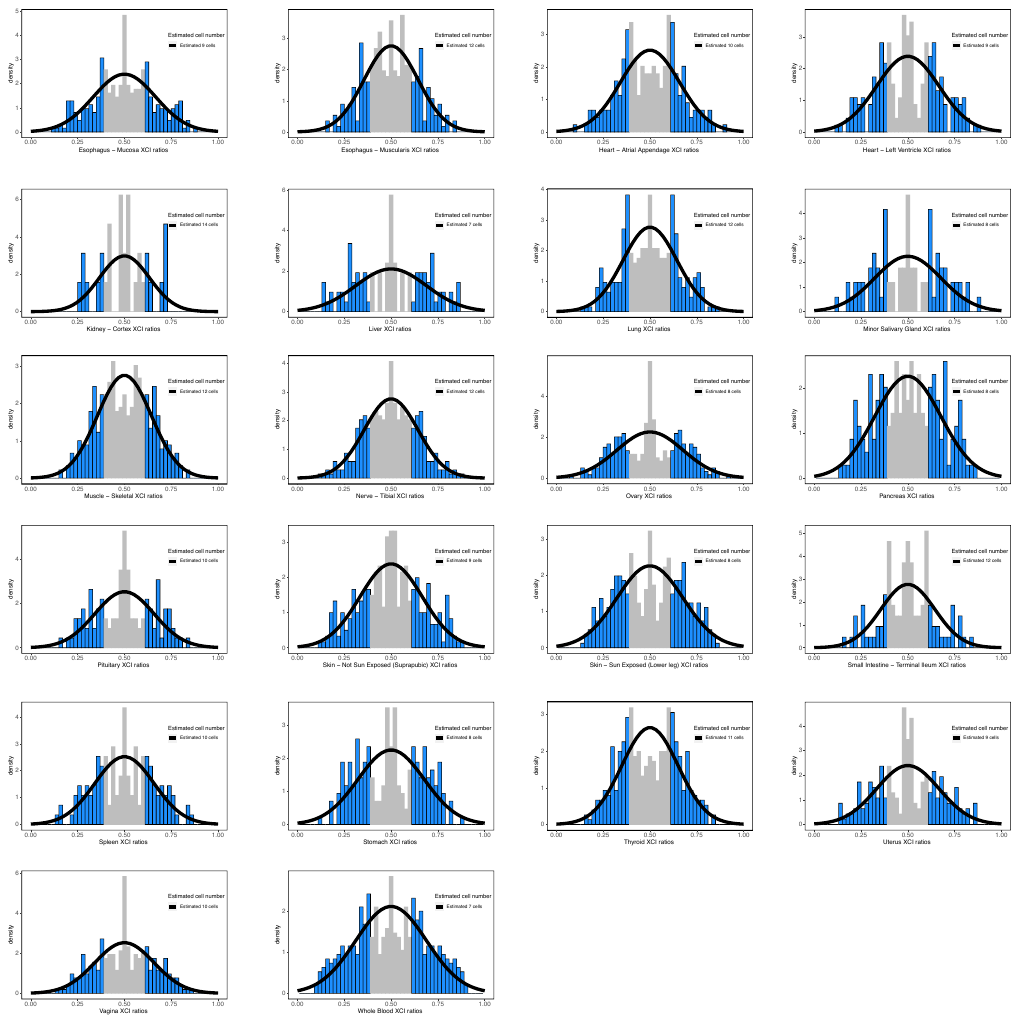

Variance in population XCI ratios maps to a pool of 6-16 cells present at the time of XCI completion

**a**, Histograms of tissue-specific distributions of XCI ratios. The fitted normal distribution to the tails of the empirical XCI ratio distribution (blue) is shown in black. The uncertain 0.4 – 0.6 XCI ratio estimates are shaded in grey.
