## Supplementary figures and images for "Cross-tissue analysis of allelic X-chromosome inactivation ratios resolves features of human development"

### Supplemental Figure 1

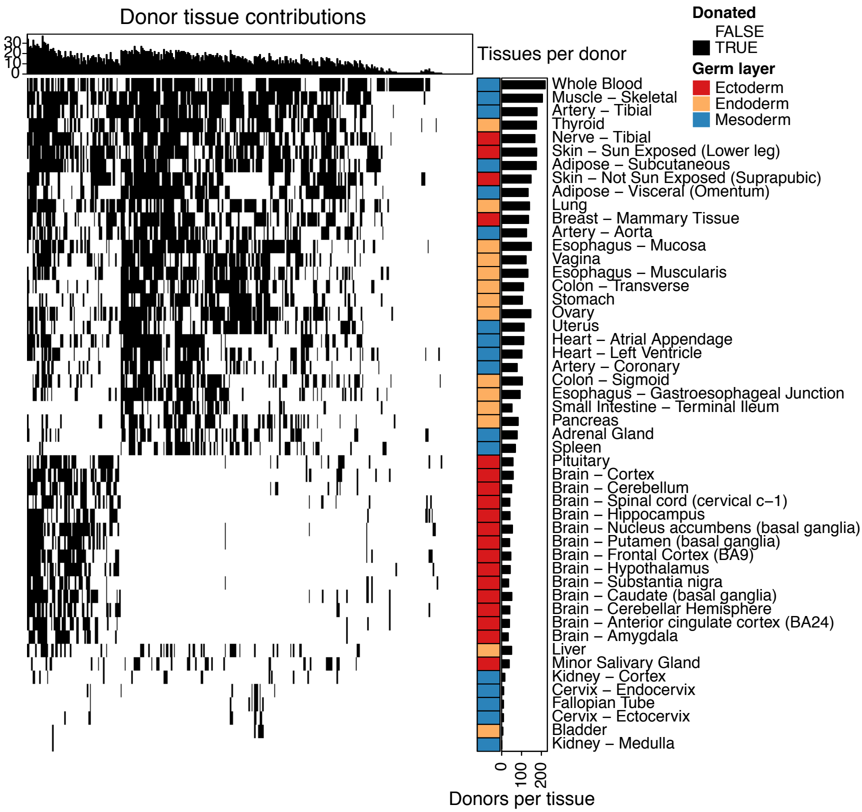

### Supplemental Figure 2

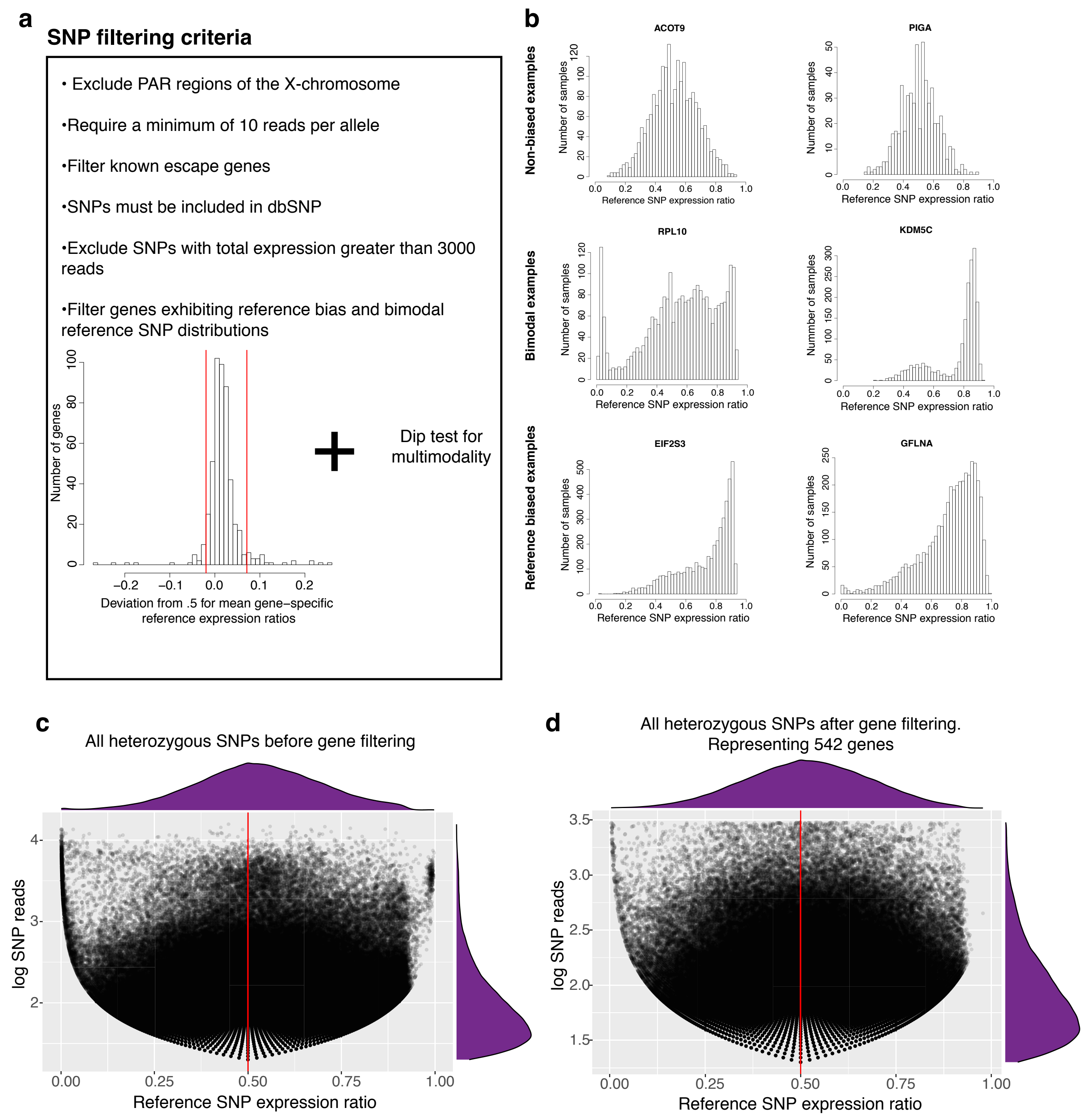

### Supplemental Figure 3

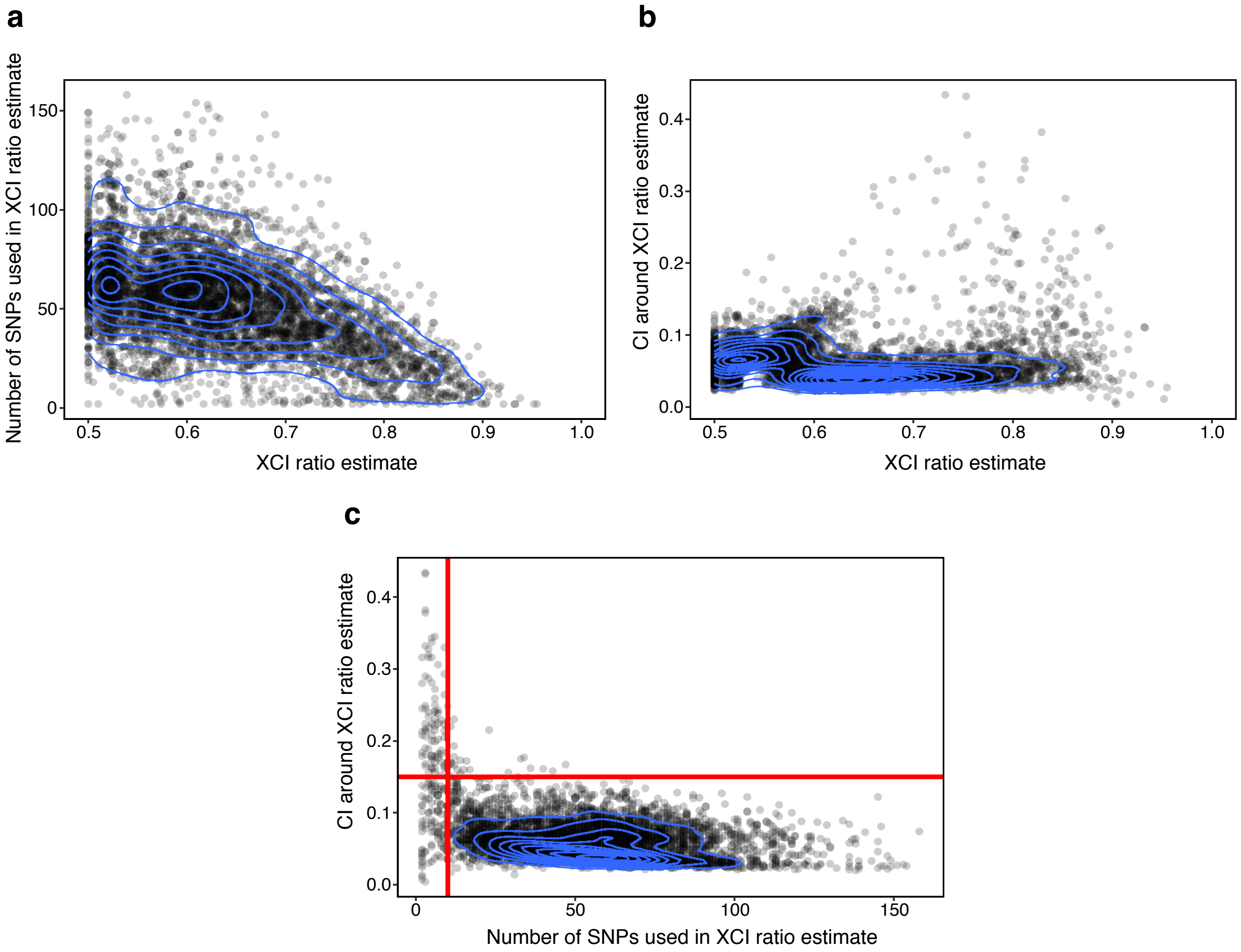

### Supplemental Figure 4

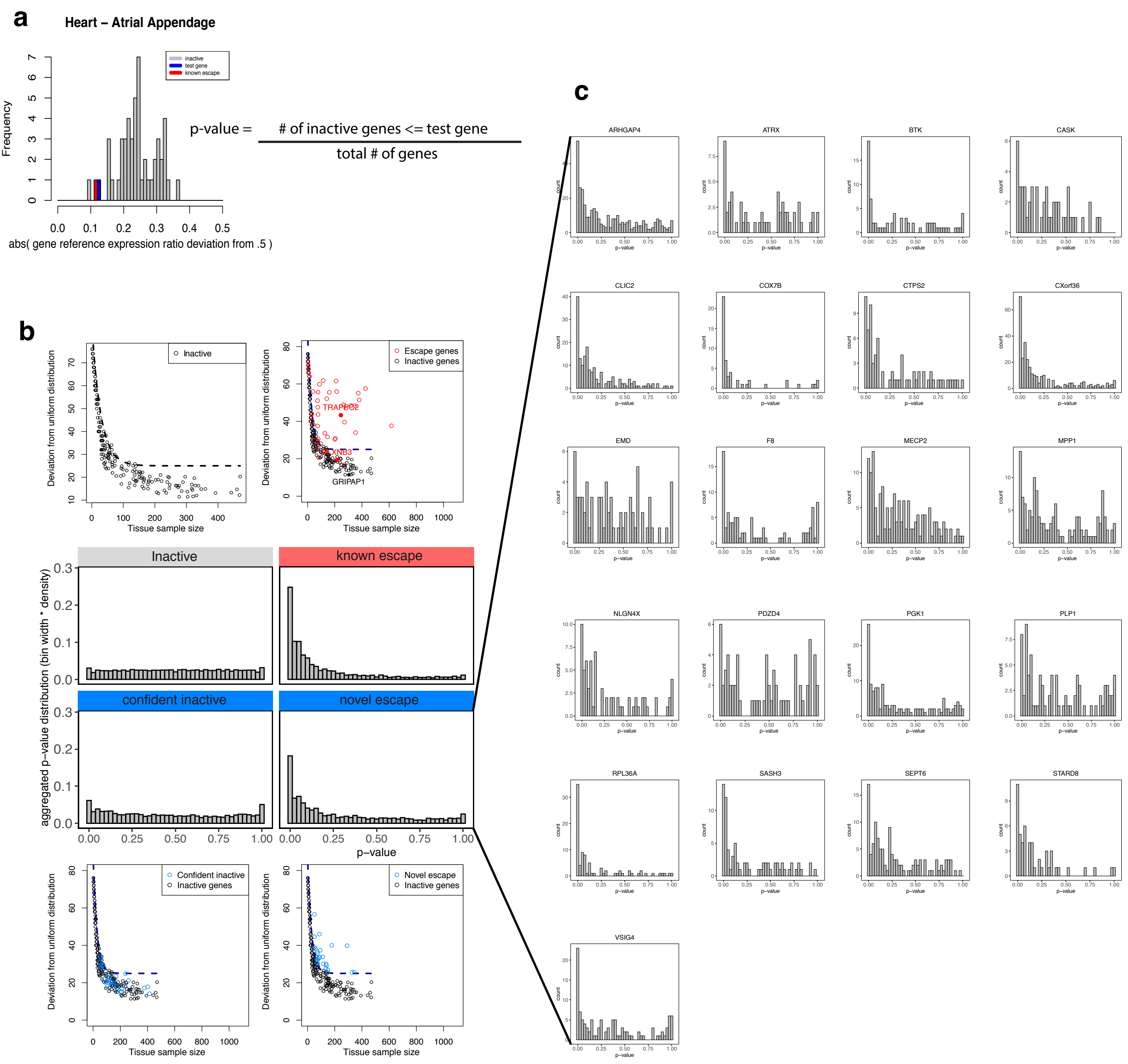

### Supplemental Figure 5

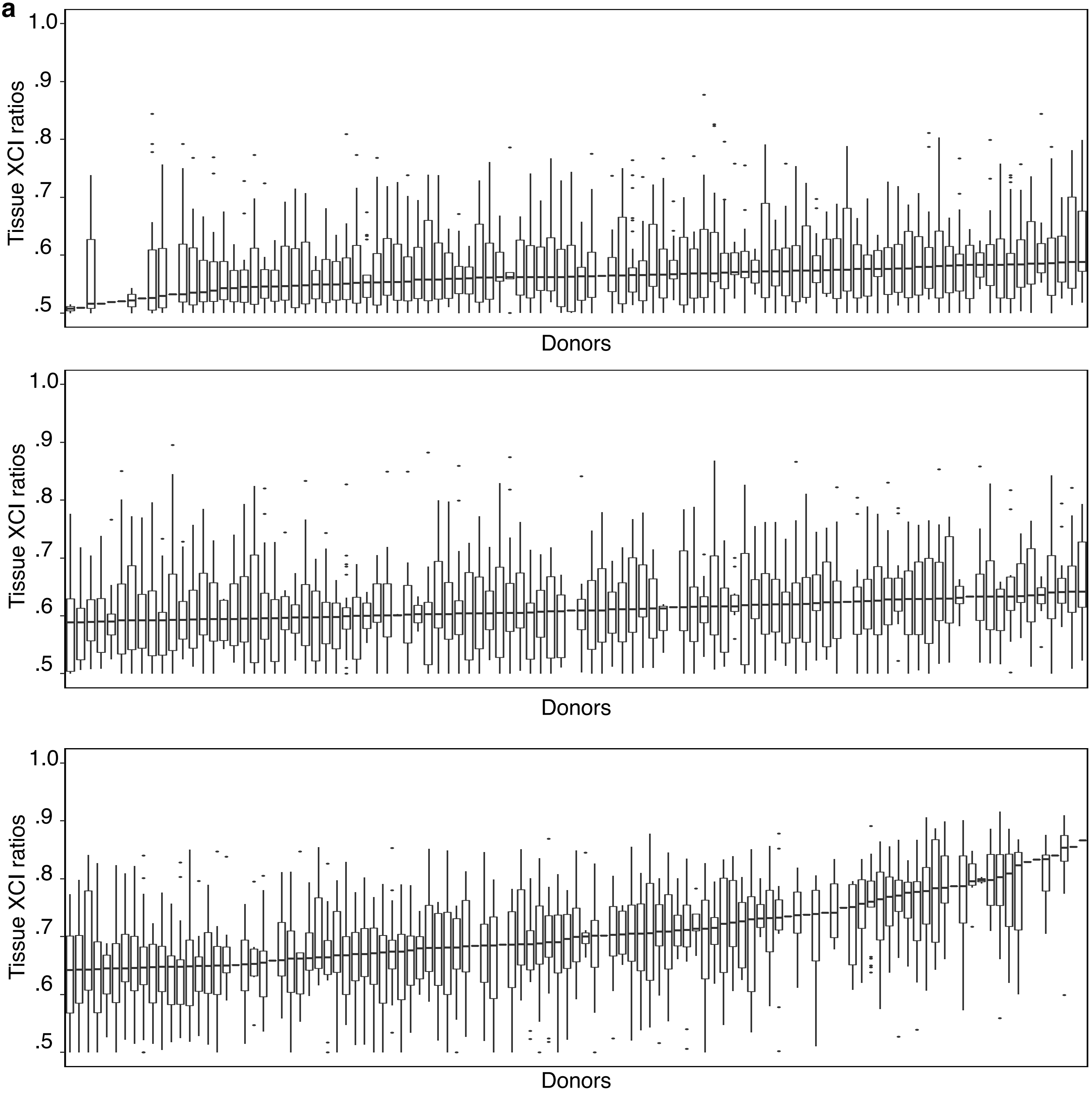

### Supplemental Figure 6

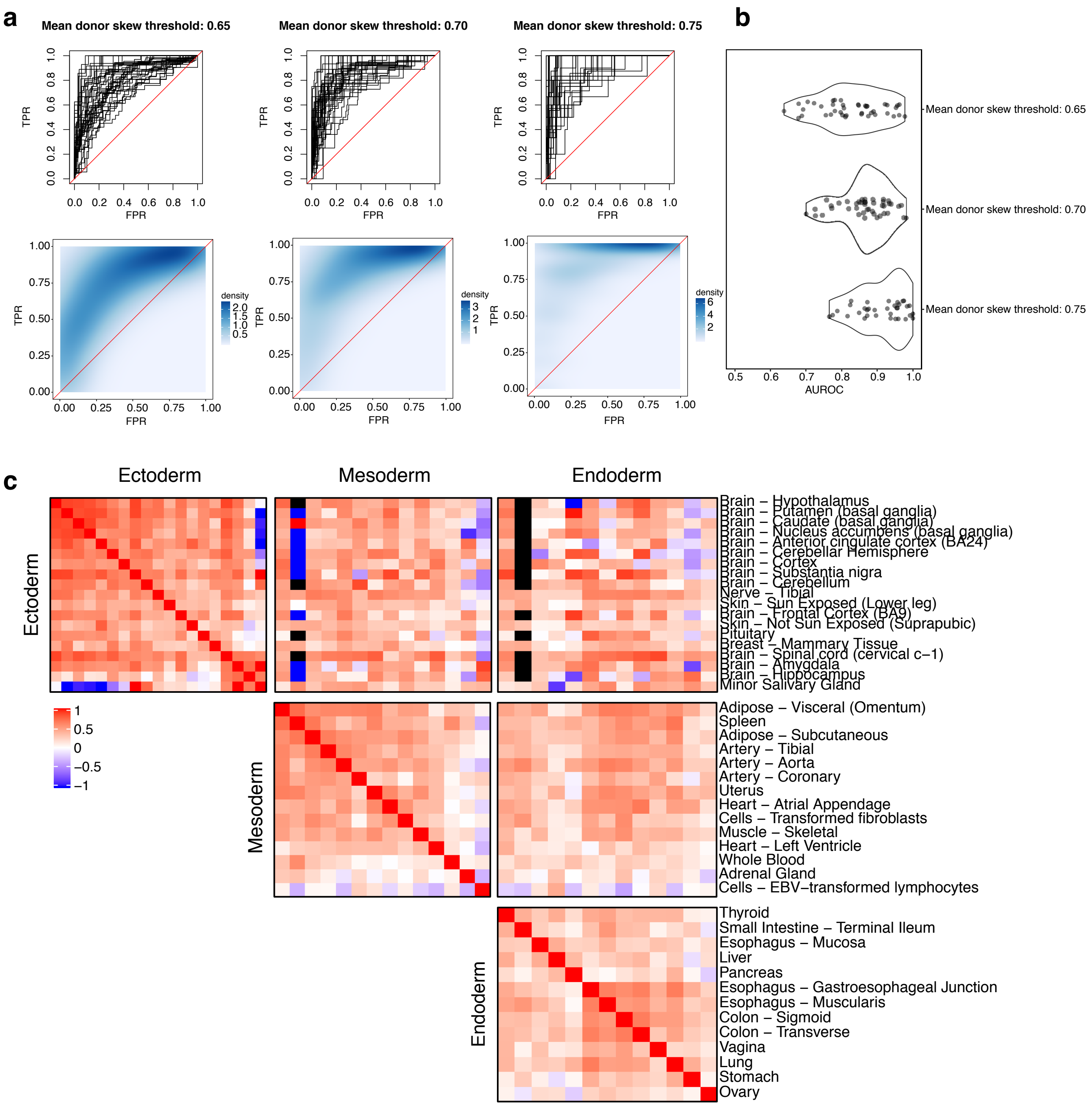

### Supplemental Figure 7

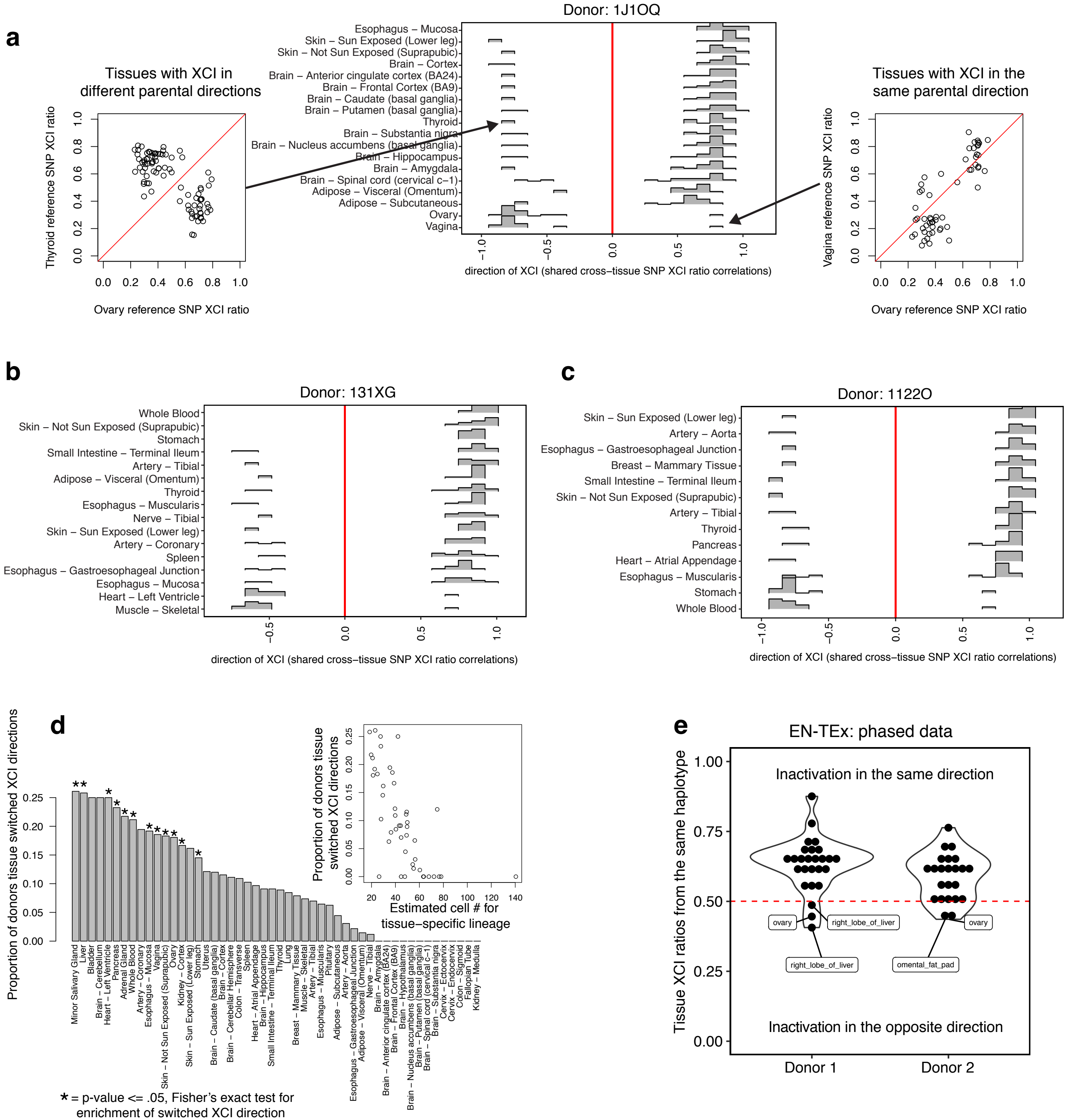

### Supplemental Figure 8.1

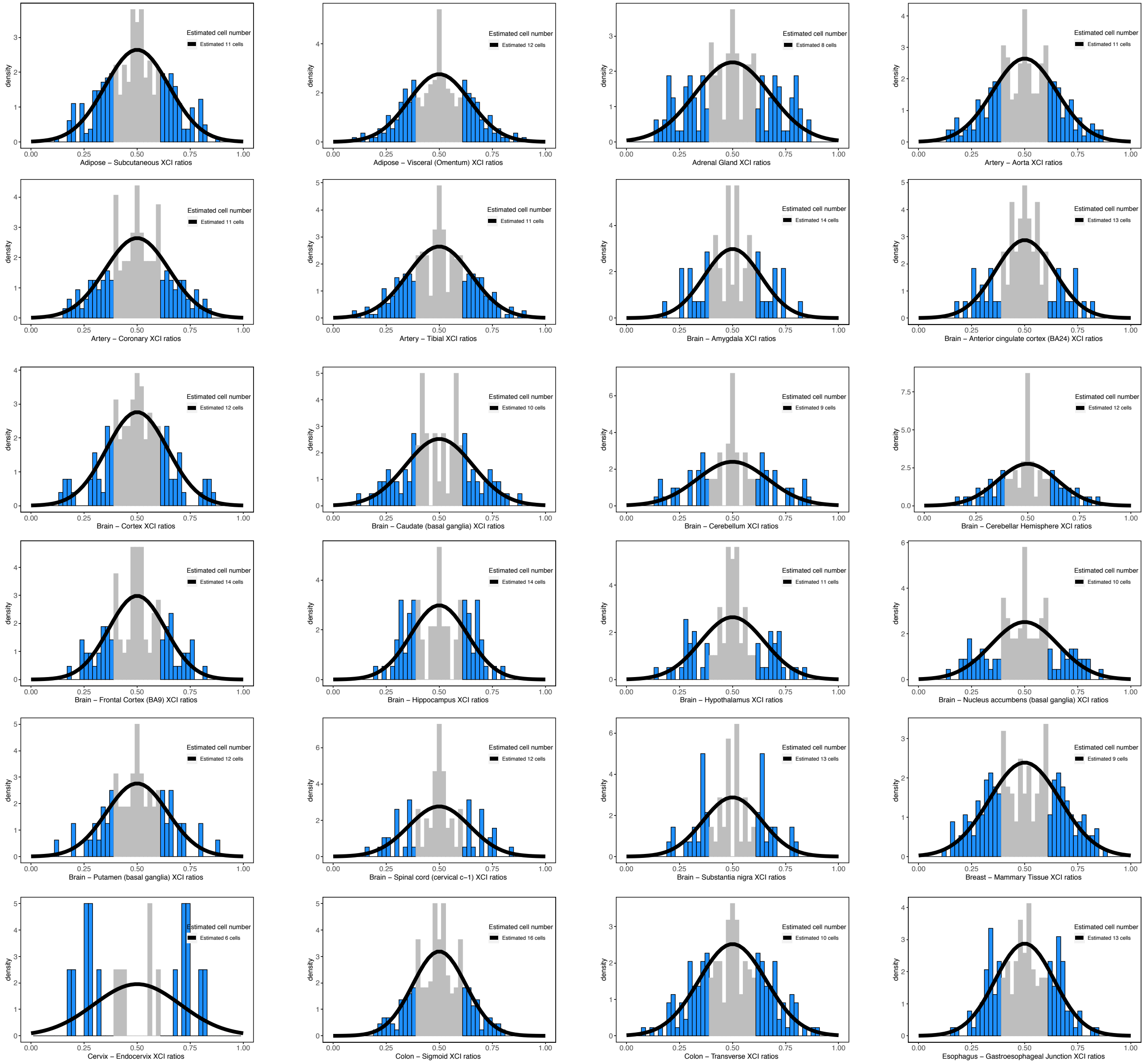

### Supplemental Figure 8.2

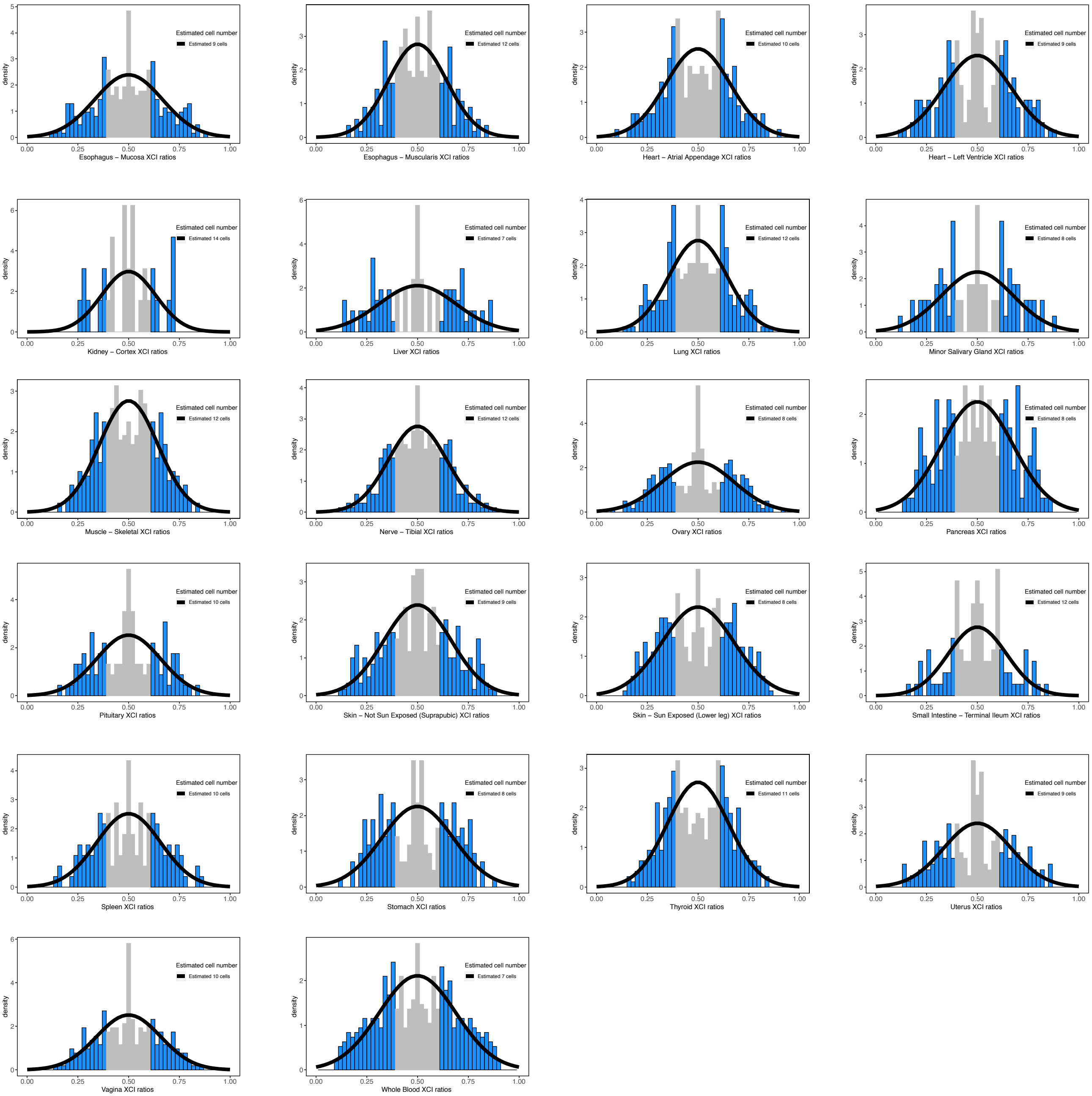
