## Supplemental Table 1 for "Cross-tissue analysis of allelic X-chromosome inactivation ratios resolves features of human development"

| gene | escape_label |
| --- | --- |
| OPHN1 | confident inactive |
| MAMLD1 | confident inactive |
| RP11-1148L6.5 | confident inactive |
| CXorf40B | confident inactive |
| WDR45 | confident inactive |
| TREX2 | confident inactive |
| ACOT9 | confident inactive |
| ARMCX5-GPRASP2 | confident inactive |
| PIR | confident inactive |
| PLXNA3 | confident inactive |
| GPC4 | confident inactive |
| DMD | confident inactive |
| HEPH | confident inactive |
| PRPS2 | confident inactive |
| SRPK3 | confident inactive |
| SLC6A8 | confident inactive |
| TMEM47 | confident inactive |
| EFNB1 | confident inactive |
| MIR503HG | confident inactive |
| PNMA3 | confident inactive |
| RP5-972B16.2 | confident inactive |
| CHRDL1 | confident inactive |
| SAT1 | confident inactive |
| USP11 | confident inactive |
| COL4A6 | confident inactive |
| TCEAL2 | confident inactive |
| TSPYL2 | confident inactive |
| PLS3 | confident inactive |
| BEX1 | confident inactive |
| TSC22D3 | confident inactive |
| MSN | confident inactive |
| SYN1 | confident inactive |
| TMSB4X | confident inactive |
| YIPF6 | inactive |
| RLIM | inactive |
| TBC1D25 | inactive |
| ZBTB33 | inactive |
| PHF6 | inactive |
| ZNF75D | inactive |
| THOC2 | inactive |
| PHKA2 | inactive |
| MORC4 | inactive |
| GAB3 | inactive |
| MTMR1 | inactive |

| | |
|---|---|
| RPS6KA3 | inactive |
| MED12 | inactive |
| INTS6L | inactive |
| SCML1 | inactive |
| WDR13 | inactive |
| APEX2 | inactive |
| ATP11C | inactive |
| ARHGAP6 | inactive |
| ELF4 | inactive |
| PHF8 | inactive |
| TAF9B | inactive |
| MAP7D3 | inactive |
| FAM122B | inactive |
| SLC25A43 | inactive |
| SMIM10 | inactive |
| ABCD1 | inactive |
| RBBP7 | inactive |
| DLG3 | inactive |
| C1GALT1C1 | inactive |
| SLC25A53 | inactive |
| LINC01278 | inactive |
| TBL1X | inactive |
| ZNF275 | inactive |
| TMEM164 | inactive |
| PRPS1 | inactive |
| ARMCX1 | inactive |
| FMR1 | inactive |
| HUWE1 | inactive |
| ZMAT1 | inactive |
| AIFM1 | inactive |
| SLC9A6 | inactive |
| PJA1 | inactive |
| HCFC1 | inactive |
| NSDHL | inactive |
| FAM127C | inactive |
| OCRL | inactive |
| GRIPAP1 | inactive |
| RBM10 | inactive |
| APOO | inactive |
| ZDHHC9 | inactive |
| ARMCX3 | inactive |
| ZMYM3 | inactive |
| UBL4A | inactive |
| GS1-358P8.4 | inactive |
| EBP | inactive |

| | |
|---|---|
| REPS2 | inactive |
| CLCN4 | inactive |
| WWC3 | inactive |
| LAGE3 | inactive |
| DOCK11 | inactive |
| DKC1 | inactive |
| ARMCX2 | inactive |
| SLC9A7 | inactive |
| IL13RA1 | inactive |
| VBP1 | inactive |
| MAGEH1 | inactive |
| TAZ | inactive |
| TSPAN6 | inactive |
| HDAC6 | inactive |
| SNX12 | inactive |
| GABRE | inactive |
| TFE3 | inactive |
| TSR2 | inactive |
| HTATSF1 | inactive |
| TIMM17B | inactive |
| LINC01420 | inactive |
| FAM3A | inactive |
| RBMX | inactive |
| PDHA1 | inactive |
| FAM127B | inactive |
| XIST | inactive |
| G6PD | inactive |
| PRAF2 | inactive |
| CD99L2 | inactive |
| BEX4 | inactive |
| HNRNPH2 | inactive |
| NONO | inactive |
| LDOC1 | inactive |
| ATP6AP2 | inactive |
| TCEAL3 | inactive |
| MAGED1 | inactive |
| MAOB | inactive |
| SH3BGRL | inactive |
| GDI1 | inactive |
| IDS | inactive |
| ATP6AP1 | inactive |
| MORF4L2 | inactive |
| RBM3 | inactive |
| MAGED2 | inactive |
| TSIX | inactive |

| | |
|---|---|
| TCEAL4 | inactive |
| SLC25A5 | inactive |
| ARSD | known escape |
| CA5B | known escape |
| CDK16 | known escape |
| CXorf38 | known escape |
| DDX3X | known escape |
| EIF1AX | known escape |
| GEMIN8 | known escape |
| GPM6B | known escape |
| GYG2 | known escape |
| IQSEC2 | known escape |
| KDM6A | known escape |
| MAOA | known escape |
| MED14 | known escape |
| MSL3 | known escape |
| MXRA5 | known escape |
| NAP1L3 | known escape |
| PLXNB3 | known escape |
| PNPLA4 | known escape |
| PRKX | known escape |
| RENBP | known escape |
| RP11-706O15.1 | known escape |
| SMC1A | known escape |
| STS | known escape |
| SYAP1 | known escape |
| SYTL4 | known escape |
| TRAPPC2 | known escape |
| TXLNG | known escape |
| UBA1 | known escape |
| USP9X | known escape |
| XG | known escape |
| ZFX | known escape |
| ZRSR2 | known escape |
| CTPS2 | novel escape |
| CASK | novel escape |
| ATRX | novel escape |
| STARD8 | novel escape |
| NLGN4X | novel escape |
| MECP2 | novel escape |
| CLIC2 | novel escape |
| F8 | novel escape |
| PGK1 | novel escape |
| SEPT6 | novel escape |
| MPP1 | novel escape |

| | |
|---|---|
| EMD | novel escape |
| CXorf36 | novel escape |
| ARHGAP4 | novel escape |
| BTK | novel escape |
| COX7B | novel escape |
| RPL36A | novel escape |
| VSIG4 | novel escape |
| SASH3 | novel escape |
| PDZD4 | novel escape |
| PLP1 | novel escape |
